# Deep learning predicts haematopoietic stem cell ageing from 3D chromatin images

**DOI:** 10.64898/2025.12.11.693143

**Authors:** Pablo Iáñez Picazo, Eva Mejía-Ramírez, Dario Di Bari, Elena Vitali, Maria Carolina Florian, Paula Petrone

## Abstract

The functional decline of the haematopoietic system during ageing propagates detrimental effects on the whole organism, ultimately eroding life and healthspan. Quantifying haematopoietic ageing holds great scientific and clinical relevance. Alterations in chromatin architecture are a well-established hallmark of ageing that encode rich and informative signatures of the ageing process, yet they remain largely unexplored as quantitative markers. Here, we present an interpretable deep learning approach based on convolutional neural networks, ChromAgeNet, that learns changes in the spatial features of chromatin architecture during natural aging of Hematopoietic Stem Cells (HSCs). We trained our algorithm on 3D microscope images of DAPI-stained HSC nuclei to discriminate between young and aged murine HSCs, achieving and AUROC of 0.77 ± 0.03. This approach outperforms classical machine learning models trained on handcrafted chromatin features from the same dataset. We then applied explainable artificial intelligence techniques, identifying chromatin entropy, peripheral heterochromatin and chromatin condensates as predictive markers. As a proof of concept, we evaluated the potential of our model as a phenotypic screening tool for aged HSCs treated with epigenetic drugs to detect rejuvenation. Altogether, we demonstrate that changes in chromatin organization can be modeled via machine learning to predict cellular ageing in the hematopoietic compartment. Our developed framework, ChromAgeNet, serves as an interpretable algorithm to unravel the intricate relationship between chromatin changes and cellular ageing, and advance high throughput drug screening for rejuvenation therapies.

## 1 Background

Age is the leading risk factor for prevalent human diseases in developed countries [1]. Ageing, defined as time-related biological deterioration, originates fundamentally at the cellular and subcellular levels. The hematopoietic system is particularly compromised during aging, which affects all of its components [2]. At the cornerstone of this system lie the Haematopoietic Stem Cells (HSCs), which play a crucial role in maintaining lifelong blood and immune homeostasis. These cells suffer from a functional decline upon ageing that drives the impairment of the haematopoietic system in the elderly [3, 4]. A collection of twelve general hallmarks of cellular ageing has been proposed, including cellular senescence, stem cell exhaustion and epigenetic alterations [5], yet the precise processes driving HSC functional deterioration upon ageing are not fully understood [6]. There is a growing need for approaches that can predict the cellular age of individual HSCs and decipher the molecular features that distinguish youthful from aged stem cells.

Epigenetic clocks based on DNA methylation have been shown to be powerful tools for estimating biological age [7], inspired by the pioneering works of Hannum [8] and Horvath [9], and are being used to predict mortality and disease risk, evaluate rejuvenation therapies, and identify ageing-related factors [10]. But other cellular hallmarks of ageing, including telomere length [11], genomic instability [12], and cellular senescence [13], can be used to construct age prediction models [14]. In addition, emerging high-dimensional molecular readouts, such as transcriptomics [15, 16], proteomics [17, 18], metabolomics [19, 20], and multiomics [21] can capture complementary dimensions of ageing. Notably, the nuclear architecture and spatial organization of chromatin within the cell nucleus also deteriorate over time [22] and offer an untapped but rich source of information that can be modeled for age measurement.

Chromatin conformation studies have revealed that the genome is intricately organized in space, spanning a multiscale hierarchy of chromatin condensation. It includes chromosome territories, A/B compartments, topologically associating domains (TADs), and chromatin loops [23–25]. This 3D nuclear architecture is altered at all scales during ageing and senescence [25], resulting in the loss of facultative heterochromatin [26–29], redistribution of chromatin from the nuclear periphery toward the interior [29–31], and changes in histone modifications [32]. An example of this is the erosion of Lamin-Associated Domains (LADs) of the chromatin, resulting in the detachment of peripheral constitutive heterochromatin from the nuclear lamina [31]. As for HSCs, we previously reported that decreased expression of Lamin A/C contributes to altered nuclear shape and volume in aged HSCs [33]. More recently, we further characterized mechanical alterations and the changes in nuclear envelope folding and tension in aged HSCs [34]. New advances in confocal microscopy and imaging techniques have progressively improved the capacity to visualize and quantify structural changes of the nucleus [35], opening the door to image-based models of nuclear ageing.

Texture features of the nucleus have long been employed to describe cellular heterogeneity and categorize disease states in microscopy images [36, 37]. Machine learning (ML) approaches can be trained with these handcrafted features to examine chromatin organization [38, 39]. More recently, image-derived endogenous histone modifications have been used to measure drug response [40] and estimate epigenetic age [41]. The revolutionary emergence of deep learning (DL), and particularly convolutional neural networks (CNNs) [42], has enabled models to learn directly from raw image data, bypassing the need to predefine a set of features, and achieving state-of-the-art performance in nuclear classification tasks. CNN algorithms have revealed that nuclear morphology is a robust predictor of cellular senescence across imaging modalities including confocal, phase contrast, and H&E-stained images [43–46]. More recently, [47] trained a CNN on images of histone modifications and RNA Pol II to predict phenotypic heterogeneity, identifying the nucleolus as an important discriminative marker. While CNNs have also been applied for HSC and Multipotent Progenitors (MPPs) classification [48] and drug screening [49, 50], to the best of our knowledge, no prior work has modeled chromatin structure at subnuclear resolution in 3D images during physiological stem cell ageing.

Here, we build DL models trained on a dataset of high-resolution 3D DAPI-stained confocal images to investigate spatial changes in the nuclear architecture of single HSCs during aging. To this end, we develop **ChromAgeNet**, a CNN-based model trained to predict the biological age of HSCs. We benchmark ChromAgeNet against classical ML models trained on handcrafted chromatin features and then apply explainable AI (XAI) techniques to interpret the chromatin-associated patterns underlying model predictions. Finally, we evaluate ChromAgeNet as a proof-of-concept phenotypic screening tool by testing whether it detects rejuvenation signatures in drug-treated aged HSCs. Together, this work establishes a novel, interpretable, and scalable framework for modeling stem cell ageing and rejuvenation, demonstrating that age-related changes in the spatial arrangement of the chromatin can be computationally modeled to build algorithms for age prediction.

## 2 Results

Aged HSCs present with functional impairments that drive ageing of the whole haematopoietic system. However, HSCs are very heterogeneous and approaches that can accurately predict the cellular age of individual HSCs to distinguish youthful from aged stem cells are still lacking. Epigenetic changes that alter the chromatin architecture are suggested to contribute to the ageing of HSCs. Therefore, we explored whether chromatin architecture encoded in 3D confocal microscopy images of intact HSC nuclei stained with DAPI would be able to distinguish young from old HSCs. To this end, we developed ChromAgeNet, a CNN-based framework to classify 3D images of HSC nuclei to predict age (**Fig. 1a**).

**Figure 1.**
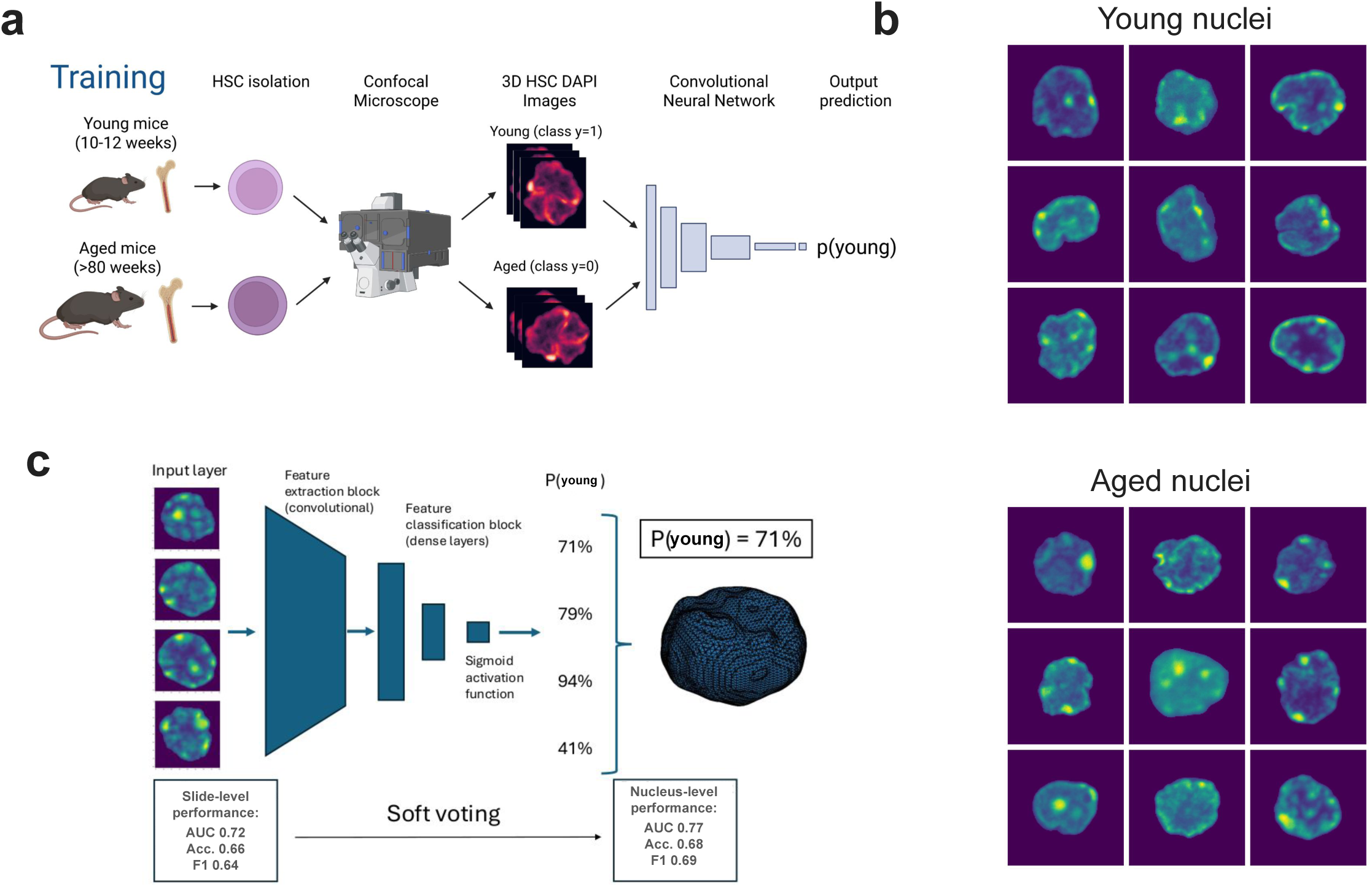
Workflow for model training and score aggregation across the 3D stack. **a.** Schematic representation of the training pipeline used to develop and train ChromAgeNet. Designed with BioRender. **b.** Representative 2D images of the XY plane for aged and young DAPI-stained HSC nuclei. These matrices are expanded using data augmentation techniques and used as input samples for ChromAgeNet. **c.** Schematic representation of the soft voting approach to aggregate ChromAgeNet probability scores obtained for each 2D nucleus slide into a whole nucleus score.

### A novel 3D dataset of DAPI-stained nuclei during HSC ageing

We collected 1,229 3D confocal microscopy images of HSC nuclei stained with DAPI to target nuclear chromatin, acquired from young (0–16 weeks old) and aged (¿80 weeks) murine BMs. To maximize training samples and decrease learning complexity, we extracted 2D slices from segmented 3D stacks (XY planes; **Supp. Fig. 1a**) of images resized to 0.05*µ*m resolution in all dimensions, applying data augmentations but preserving the nucleus size as an informative feature of ageing [22, 34]. We leveraged image interpolation on the Z axis to generate new synthetic 2D images preserving biologically-meaningful intensity distributions. Given the higher phenotypic heterogeneity expected in aged HSCs and lower ageing variability in young counterparts, we labeled young HSC images as the positive class to predict (*y* = 1), and old HSC as *y* = 0. The model was therefore trained on a balanced dataset with a total of 81,643 2D slices, comprising 36,275 slices (44.4%) from 551 young HSCs and 45,368 slices (55.6%) from 678 aged HSCs (**Fig. 1b, Table 1 and Table 2**), from which 126 nuclei were included in DMSO medium. After training, the model outputs a calibrated “youthful score” indicating the likelihood that an observed chromatin image belongs to a young HSC. The probability scores for all images belonging to the same nucleus are then aggregated via a soft voting scheme to output a single nucleus-level prediction, enhancing classification robustness across the full 3D stack (**Fig. 1c**).

**Table 1.** Summary of numerical metadata variables of original 3D images.

| Variable | Young HSCs (n = 551) | Aged HSCs (n = 552) |
| --- | --- | --- |
| **Z resolution ($\mu\text{m}$)** | $0.32 \pm 0.10$ | $0.32 \pm 0.09$ |
| **Number of Z-stacks** | $29 \pm 11$ | $28 \pm 10$ |
| **Sigma estimated noise** | $2.6 \pm 1.6$ | $2.8 \pm 1.3$ |
| **Total pixel intensity** | $2.1e^{08} \pm 1.3e^{08}$ | $2.5e^{08} \pm 1.6e^{08}$ |
| **Mean pixel intensity** | $7.4 \pm 5.3$ | $8.3 \pm 4.2$ |
| **Maximum pixel intensity** | $189 \pm 63$ | $211 \pm 55$ |

**Table 2.** Summary of categorical metadata variables of original 3D images.

| Variable | Value | Young HSCs<br>(n=551) | Aged HSCs<br>(n=552) |
| --- | --- | --- | --- |
| Acquired by | Microscopist 1 | 377 | 376 |
|  | Microscopist 2 | 52 | 175 |
|  | Microscopist 3 | 123 | 0 |
| Number of channels | 1 | 188 | 190 |
|  | 2 | 219 | 193 |
|  | 3 | 145 | 168 |
| Sex | F | 52 | 124 |
|  | M | 0 | 51 |
|  | unknown | 500 | 376 |
| Year of acquisition | 2019 | 29 | 39 |
|  | 2020 | 142 | 134 |
|  | 2021 | 11 | 23 |
|  | 2022 | 23 | 29 |
|  | 2023 | 66 | 80 |
|  | 2024 | 281 | 246 |

### Deep learning discriminates young and aged HSCs

We trained and validated our model via five cross-validation, splitting the data at the 3D nucleus level, demonstrating that ChromAgeNet achieved consistent performance across folds. The class probability histograms from a single fold revealed clear density separation, with the majority of confusion around scores near 0.5 (**Fig. 2a**). At the image level, our model reached an average validation AUROC of 0.72±0.02, with 0.66±0.03 accuracy and an F1 score of 0.64±0.01 (**Supp. Table 1**). Aggregating predictions at the nucleus level improved performance to 0.77±0.03 AUROC, 0.68±0.05 accuracy, and 0.69±0.02 F1 (**Supp. Table 2**), with the best fold reaching a validation AUROC of 0.81 **(Supp. Fig. 1b**). In general, ChromAgeNet showed a greater sensitivity for young HSCs (78% TPR) than for aged HSCs (61% TNR; **Fig. 2b**), using decision thresholds below 0.5 that improved young class separation (**Fig. 2c,d**). We examined the distribution of scores along the Z-axis, and found that prediction stability was the highest at the nuclear midsection, where the nuclear area was the largest, and decreased near the apical and basal poles **(Supp. Fig. 1c**).

**Figure 2.**
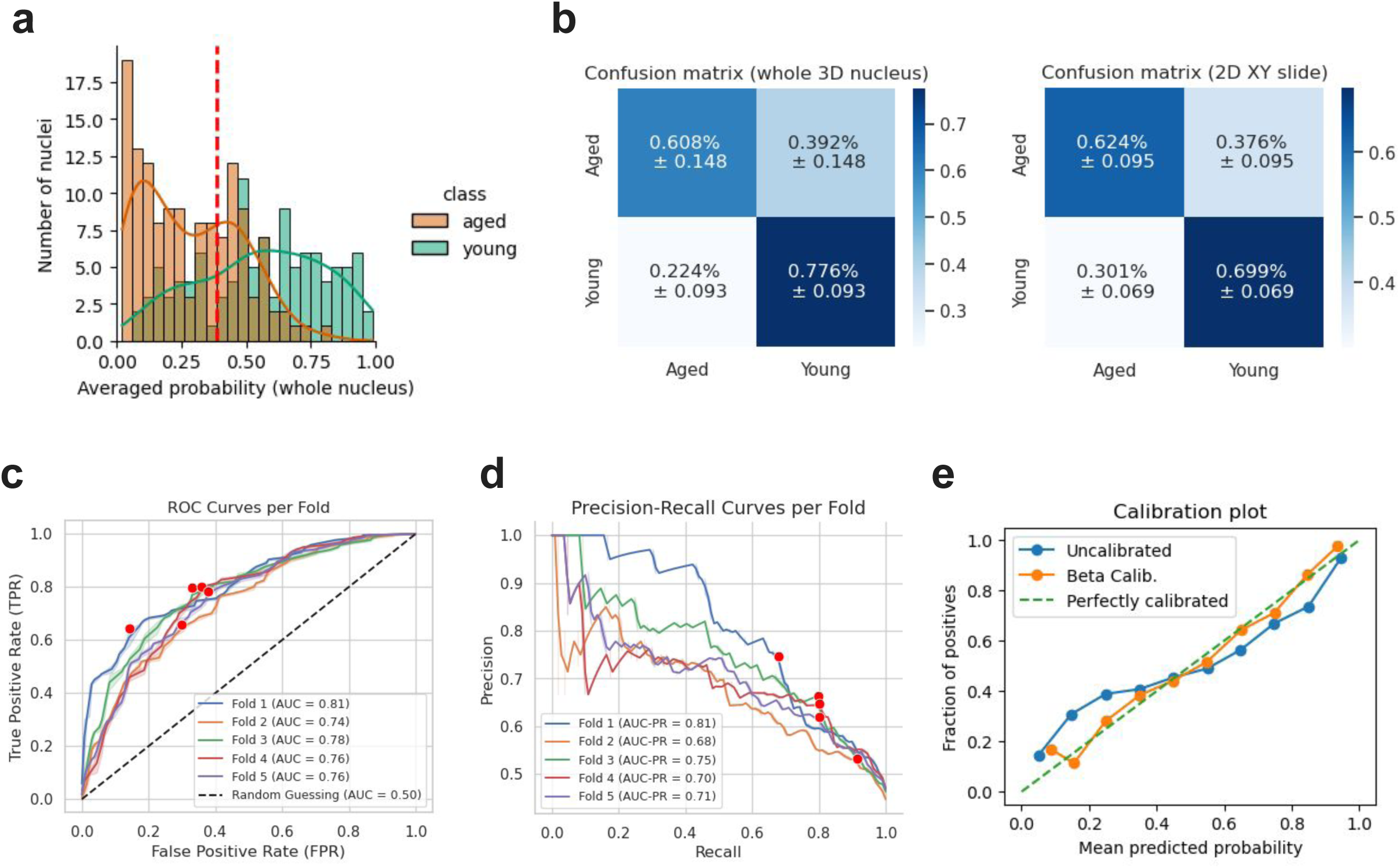
ChromAgeNet discriminates young and aged HSCs. **a.** Distributions of ChromAgeNet scores for the two ground-truth classes in the validation dataset. The curves display the kernel density estimate for each class, while the vertical red line marks the classification threshold applied to obtain predicted classes. **b.** Confusion matrices showing the performance of ChromAgeNet over the validation dataset at the whole 3D nucleus level (left) and at the 2D slide level (right). **c.** ROC curve and AUC for ChromAgeNet performance on validation data for each 5 CV data splits. The classification threshold yielding the highest AUC score is shown as a red dot. **d.** Precision-Recall curve and PRAUC for ChromAgeNet performance on validation data for each 5 CV data splits. The classification threshold yielding the highest AUC score is shown as a red dot. **e.** Calibration plot displaying the agreement between ChromAgeNet scores and the frequency of observations at different deciles for the uncalibrated model, and the beta-calibrated model. Results are shown for one CV data iteration. Probability calibration is essential for producing reliable predictions suitable for meaningful biological interpretation. A perfectly-calibrated algorithms follows the dashed diagonal line.

To ensure that ChromAgeNet’s youthful scores reflect true class likelihoods, we applied Beta Calibration [51] to align the output probabilities with observed frequencies. This procedure generated well-calibrated scores (Expected Calibration Error (ECE) = 0.02±0.01; Brier Score (BS) = 0.22±0.01) without sacrificing prediction performance (**Supp. Fig. 1d**), producing less extreme (highly confident) and more reliable probabilities (**Fig. 2e, Supp. Fig. 1e**). We therefore report calibrated scores in downstream inferences.

In general, these results suggest that age-related changes are encoded in 3D chromatin organization and they can be detected using DL algorithms trained on DAPI-stained slides to predict young versus aged HSCs.

### Comparison with models trained on chromatin-extracted features

To benchmark ChromAgeNet against traditional nucleus classification approaches, we trained ML models on a set of handcrafted chromatin features extracted from segmented 3D nuclear volumes resized to 0.1 *µ*m in all dimensions (**Fig. 3a**, and more in detail in **Supp. Fig. 2a**) on original and filter-transformed images (**Supp. Fig. 2b**). After filtering out highly correlated variables, we retained 70 features capturing intensity, texture, and shape for analysis. **(Supp. Fig. 2c, Supp. Data Table 1**).

**Figure 3.**
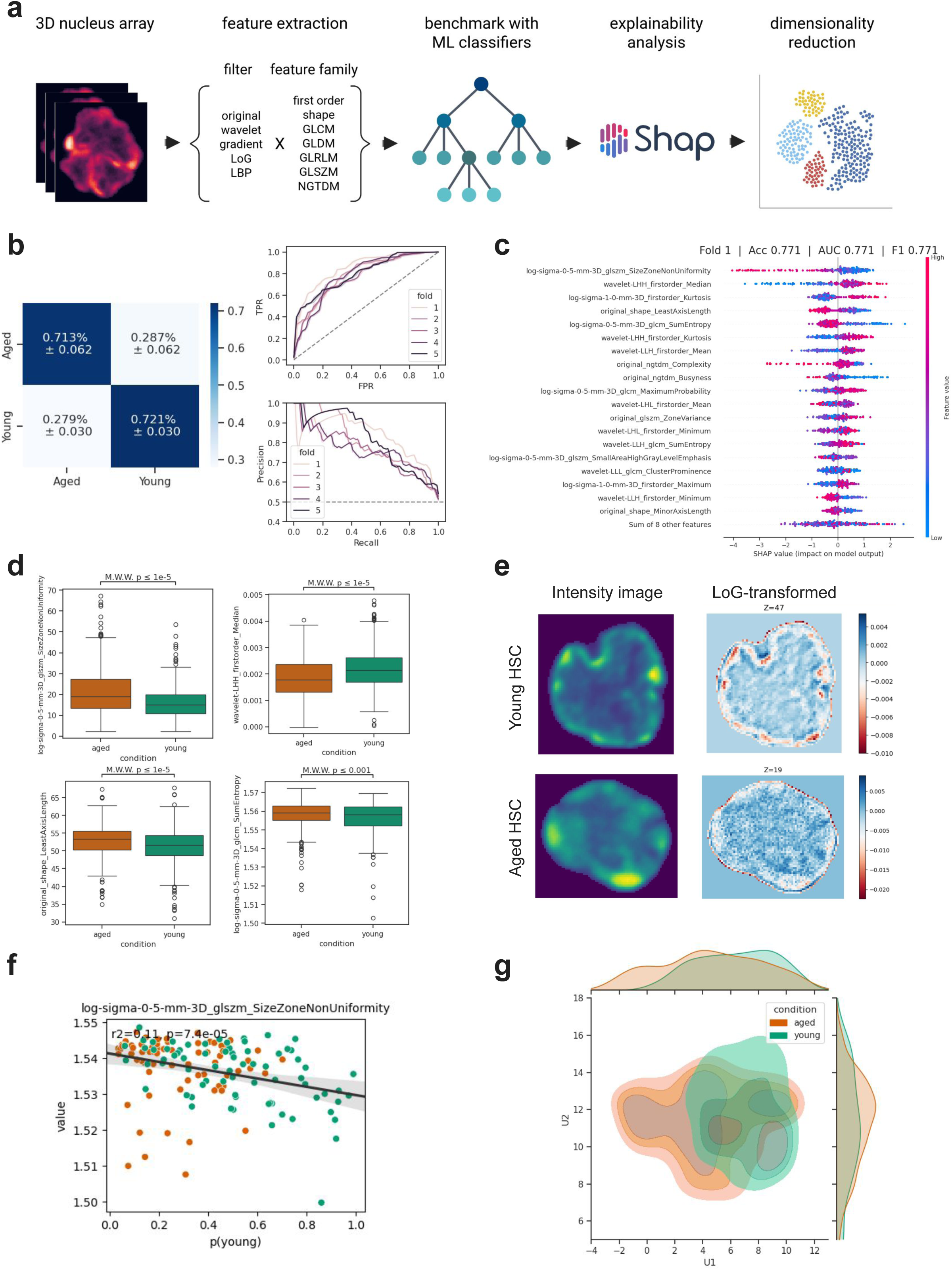
Chromatin-extracted features inform on stem cell ageing. **a.** Representation of the pipeline to extract textural and shape descriptors from 3D whole nucleus images. Designed with BioRender. **b.** Plots showing the performance of our best model, including the confusion matrix averaged over all folds, and the ROC and Precision-Recall curves for each of the data splits in our cross-validation pipeline. **c.** Summary plot of feature importance calculated using SHAP values for the first cross-validation fold. The X axis represents the impact of each model on the model’s output (contribution toward the young HSC class), while the color shows the scaled feature value (higher values are colored as red, while lower values are colored as blue). **d.** Boxplot representing differences in selected top SHAP features, X (left) and Y (right). Mann-Whitney-Wilcoxon U-test p-values with Bonferroni correction are indicated. **e.** Representative 2D images of young and aged HSC nucleus, and the resulted transformation with Laplacian of Gaussian (sigma=0.5) filter. **f.** Scatterplots showing the relationship between ChromAgeNet scores and selected top SHAP features. A regression line is shown to demonstrate the strength of correlation, along with computed *R*^2^ and p-value. **g.** UMAP embedding calculated using the top SHAP features across all CV data splits. Marginal distributions are shown along the plot edges for each component. Class densities are represented as kernel estimations to better illustrate the separation among young and aged HSCs.

We compared four shallow ML classifiers and eight feature selection techniques, resulting in 32 different modeling combinations (see Methods). A calibrated XGBoost classifier achieved the best performance (AUROC 0.73±0.04; **Fig. 3b**) in a five cross-validation setting, trained on features selected by a Random Forest (RF) model. Notably, feature selection consistently improved model performance across all tree-based classifiers **(Supp. Fig. 3a-c, Supp. Table 3**), whereas logistic regression underperformed across all feature selection scenarios **(Supp. Table 4**). The predictive performance varied modestly across folds, with AUROCs ranging between 0.67±0.02 and 0.73±0.03 (**Supp. Table 5**). Standardizing the images by binarizing intensity values resulted in slightly reduced performance metrics **(Supp. Fig. 4**).

**Figure 4.**
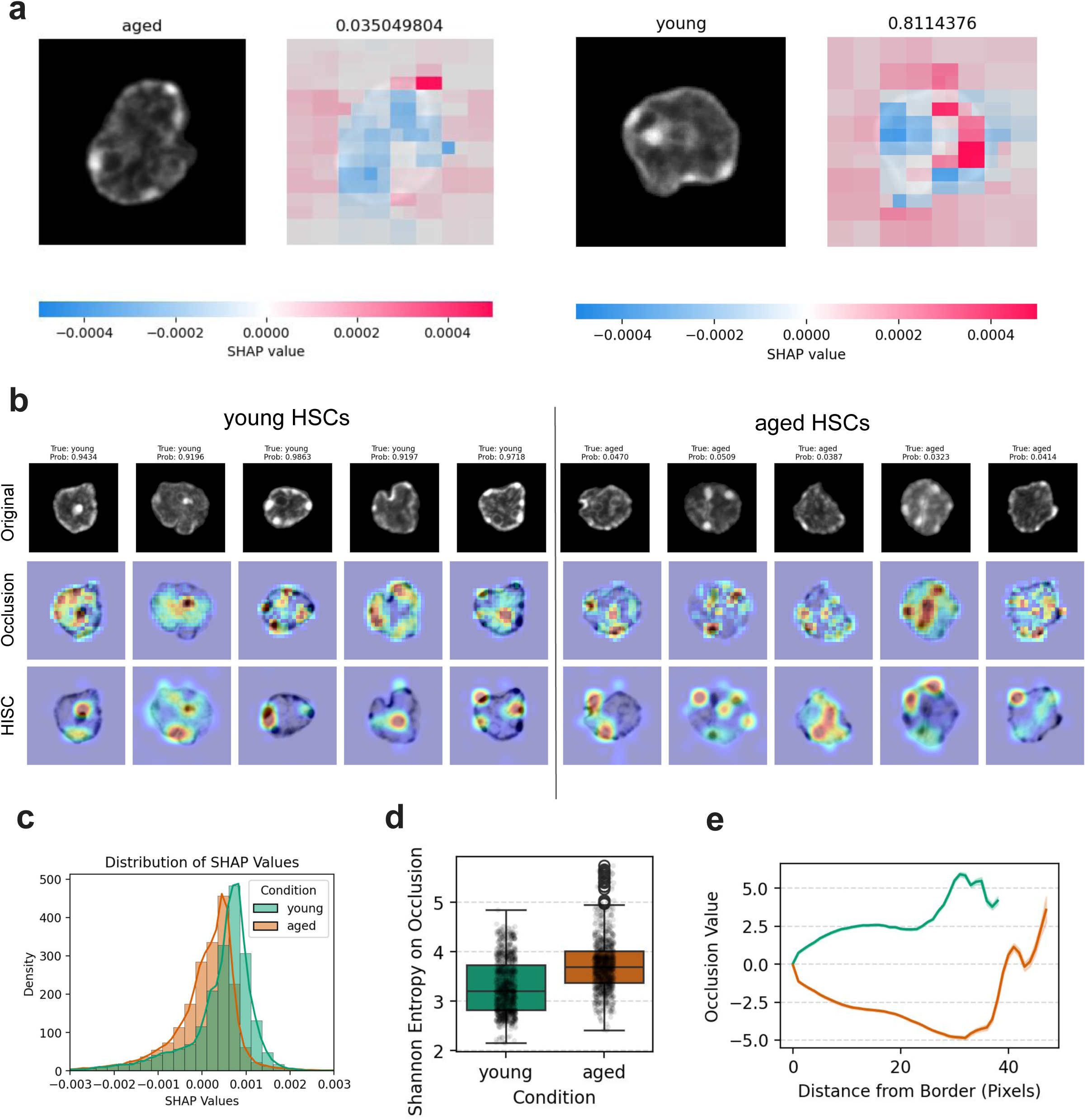
ChromAgeNet reveals spatial chromatin patterns linked to HSC ageing. **a.** Original intensity images (left) and overlaid images showing the attention maps using SHAP. Red values mean higher SHAP value, or image regions contributing the most toward the young class (with label y=1), while blue values represent lower SHAP values, or image regions contributing against the young class. A representative example for young HSC is shown on the left, and for aged HSC on the right. **b.** Comparison of XAI methods, with the top layer of images showing the original intensity images, the second layer showing attention maps from Occlusion Sensitivity, and the lower layer showing attention maps from HiSC Attribution. The ground truth and predicted ChromAgeNet score is depicted on top for each nucleus. **c.** Histograms comparing the distribution of SHAP values using 1,000 validation images for each class, colored by condition. **d.** Boxplot comparing the Shannon Entropy computed on the Occlusion attention maps at image level for both young and aged HSCs. **e.** Lineplot showing the average Occlusion values as a function of the distance to the nuclear border in number of pixels. Lines are drawn and colored separate by biological condition.

These analyses demonstrate that classical ML pipelines can also separate young and aged HSCs with significant performance, although CNNs trained on raw image data yield higher discriminative capacity.

### Chromatin-extracted features inform on stem cell ageing

To better understand which chromatin-extracted features contributed most to age classification, we computed SHAP values [52] using the best-performing model **(Supp. Data Table 2**). Across cross-validation folds, consistently top-ranked features included texture-based descriptors such as **size zone non-uniformity** and **sum entropy**, as well as **Wavelet median intensity** and **least axis length (Fig. 3c, Supp. Fig. 5**). These features showed statistically significant differences between classes (**Fig. 3d**), supporting their biological relevance. Furthermore, these differences can be observed visually in representative in transformed images, where young HSC images presented more structured patterns, less intensity variation in pixel neighborhoods and more homogeneous size zone volumes (**Fig. 3e**). In general, features from transformed images were more informative than characteristics from the original image features, especially from Laplacian of Gaussian (LoG) and Wavelet filters, presumably because of their capacity to highlight abrupt spatial variations in chromatin profiles. A greater number of textural features from LoG (*σ* = 0.5) filter appeared on top compared to LoG (*σ* = 1). The sigma (*σ*) parameter of LoG controls the spatial scale, suggesting that lower-frequency details such as small-scale nuclear texture variations are more relevant than larger-scale, coarse textures are. Furthermore, some of these top SHAP features showed a modest association with ChromAgeNet scores (**Fig. 3f**), indicating that our deep learning model captures a portion of the same variability.

**Figure 5.**
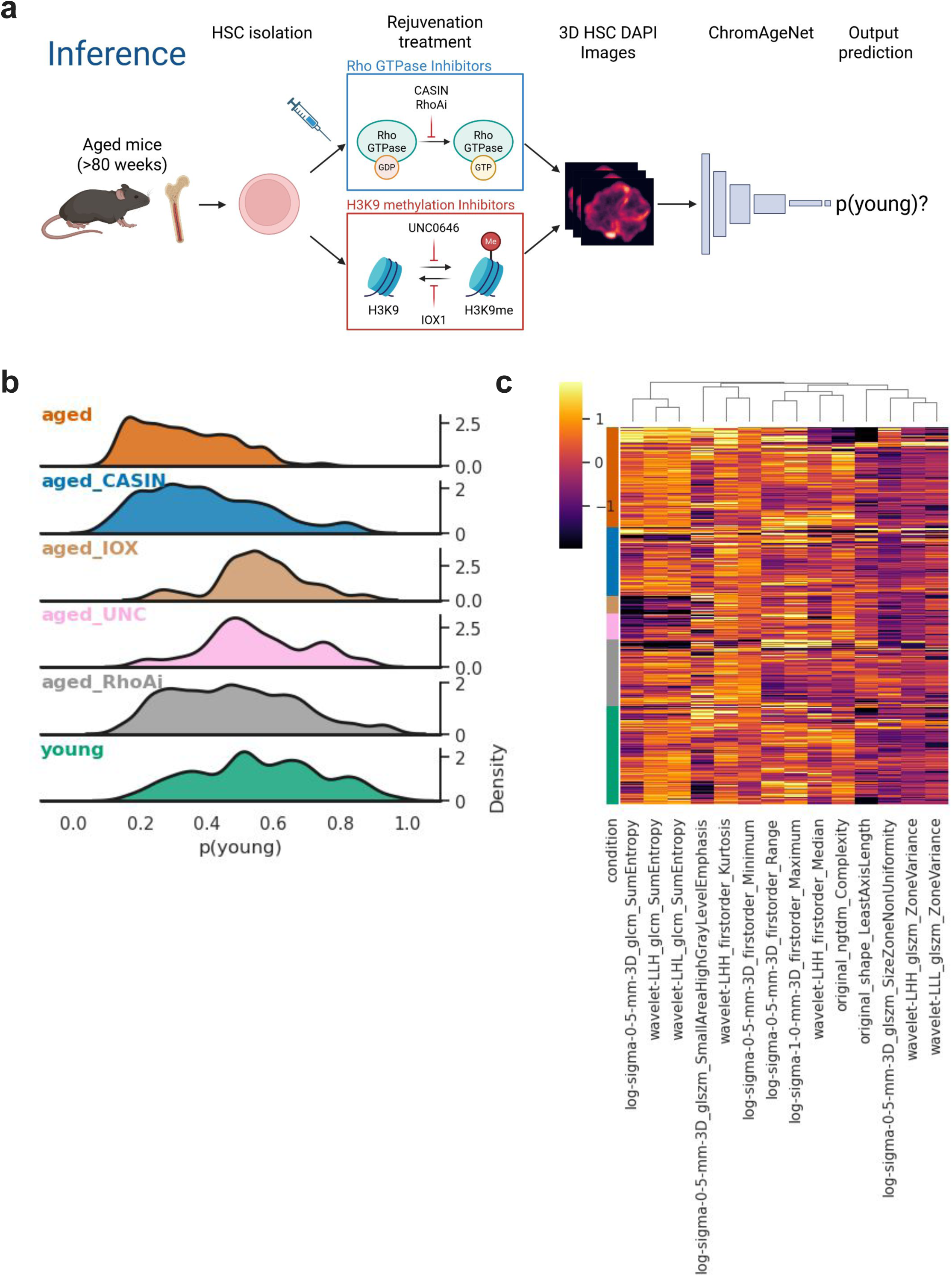
ChromAgeNet measures rejuvenation outcomes. **a.** Schematic representation of the inference pipeline using ChromAgeNet, where the two different mechanism of actions of the epigenetic drugs are depicted: Inhibitors of Rho GTPase (CASIN and RhoA inhibitor) and modulators of H3K9 methylation (UNC0646 and IOX1). Designed with BioRender. **b.** Distribution plots showing soft voting-aggregated and calibrated ChromAgeNet scores at nucleus level for young and aged HSC, along with different treatments of aged HSCs. Probability values near 0 reflect more aged-like phenotypes, while values near 1 reflect more young-like phenotypes. **c.** Heatmap showing normalized values of top selected SHAP features (columns) distributed over images from different young, aged and drug-treated aged HSCs (rows), all acquired by the same microscopist and microscope.

Using the top SHAP-ranked features across folds, we constructed a UMAP embedding that revealed a clear separation between young and aged HSCs densities (**Fig. 3g**). Altogether, while chromatin-extracted features yield lower discriminative performance than CNN-based models, they provide interpretable insights into chromatin heterogeneity and contribute to understanding nuclear changes in HSC ageing.

### ChromAgeNet reveals spatial chromatin patterns linked to HSC ageing

Despite the greater discriminative power of using ChromAgeNet over ML baselines, the internal representations of CNNs learned from the data are not readily interpretable. To uncover which chromatin patterns in our images drive model decisions, we applied a suite of explainable XAI techniques.

Using **activation maximization** [53], we visualized progressively complex patterns of chromatin signal learned by our model and associated with young HSC, as the layer depth increased in the network. This technique produced spatial chromatin textures specifically tailored for DAPI and young HSCs **(Supp. Fig. 6**). To interpret the prediction of the model at the image level we employ attribution methods, a form of local explainability that highlights regions in the pixel space that are most influential on the model output.

SHAP can also be used as an explainability method in images to highlight important pixels (features) contributing to class prediction. This technique revealed that age-associated signals contributing to both young and aged classes are present simultaneously in most images, particularly within the nuclear matrix, indicating high intra-class variability in chromatin textures (**Fig. 4a**). Recognizing that results can vary significantly between attribution methods, we systematically benchmarked a range of techniques implemented in the **Xplique** library [54] in our data. To assess the reliability of each approach, we used **MuFidelity** metric scores, which evaluate whether highlighted pixels truly affect model predictions when perturbed [55].

Among all the tested methods, **occlusion sensitivity**, a model-agnostic approach that measures the effect of masking small patches of image regions, consistently achieved the highest scores in our dataset (Methods). We compared this method with advanced gradient-based attention maps [56]. Interestingly, attention maps reveal chromatin differences between confidently predicted young and aged HSCs (**Fig. 4b**). Although nuclear morphology of aged HSCs is often more irregular, nuclear invaginations are often captured as important in young HSCs, as reported in previous studies [33]. In young HSCs, occlusion results in greater attention along the nuclear envelope (NE), whereas large and rounded DAPI blobs are typically [**?**] highlighted in aged HSCs [34]. Upon visual inspection, young HSCs tended to exhibit a continuous thin line along the NE, presumably corresponding to LAD-associated facultative heterochromatin. In contrast, aged HSCs displayed larger DAPI blobs and a thicker, irregular DAPI signal along the NE, suggestive of an age-related heterochromatin deterioration, as previously reported with other tools [33, 57]. In addition, we observed that low intensity regions surrounded by a rim of chromatin, probably corresponding to nucleoli, were frequently highlighted, particularly when adjacent to other large DAPI-dense domains (**Fig. 4b**).

We next sought to systematically quantify the spatial and statistical properties of attention maps in order to better understand what ChromAgeNet is learning. As expected, positive image SHAP values were more prominent in the young HSC class, although they were also found in aged HSC images (**Fig. 4c**). Similarly, occlusion maps generated from aged HSCs exhibited higher Shannon entropy (**Fig. 4d**), suggesting more dispersed and heterogeneous attention. Moreover, by profiling attention values using the occlusion method as a function of distance to the nuclear border as defined by our segmentation mask, we found a consistent trend of positive values for young HSCs and negative values for aged HSCs, with increased attention around the 30 pixel mark (1.5µm) from the border (**Fig. 4e**).

Overall, these analyses reveal a rich and spatial landscape of chromatin alterations associated with HSC ageing, showcasing ChromAgeNet’s ability to detect and explain these signatures in the nuclear architecture of each image.

### Deep learning detects chromatin changes in drug-treated aged HSCs

Next, we evaluated whether ChromAgeNet could detect changes in the organization of the chromatin in aged HSCs treated with candidate rejuvenating compounds. By measuring shifts in the distributions of predicted youthful scores, we quantified the extent to which drug-treated aged HSCs recovered chromatin patterns typical of young nuclei.

We tested four compounds grouped into two mechanisms of action: (1) **Rho GTPase inhibiors**, including **CASIN** (a Cdc42-specific inhibitor) [33, 58] and Rhosin, a **RhoA inhibitor** (RhoAi), previously linked to HSC epipolarity and nuclear remodeling through LaminA/C expression and nuclear envelope tension regulation [33, 34] and (2) **Inhibitors of heterochromatin Histone 3 lysine 9 (H3K9) methylation**, including UNC0646, a G9a/GLP methyltransferase inhibitor [59], and IOX1, a broad-spectrum histone demethylase inhibitor [60] (**Fig. 5a**).

Compared with baseline aged HSCs from the validation fold (mean youthful score: 0.34±0.14), treatment with CASIN and RhoAi modestly increased youthful scores (0.40±0.18 and 0.48±0.19, respectively). Notably, UNC0646 and IOX1 restored youthful scores to 0.56±0.13 and 0.55±0.16, closely matching untreated young HSCs (0.55±0.19) (**Fig. 5b, Supp. Table 6**), suggesting that alterations in H3K9 methylation may restore a youthful nuclear architecture in aged HSCs [3]. The inspection of top SHAP-ranked chromatin features that we identified previously across treatments revealed anticipated heterogeneity at the image level. Despite this, interesting patterns arise such as diminished sum entropy levels after treatment with H3K9 methylation inhibitors, comparable to profiles observed in a subpopulation of young HSC (**Fig. 5c**). These findings support the potential of ChromAgeNet as a scalable phenotypic screening readout in rejuvenation drug discovery.

## 3 Discussion

HSCs are critical for maintaining lifelong blood and immune homeostasis and, upon ageing, they drive the functional decline of the haematopoietic system in the elderly, contributing to major ageing-related diseases [3, 4]. The pronounced heterogeneity and unique cellular characteristics of HSCs highlight the clinical need for methods capable of predicting the cellular age of individual cells and potentially identifying features that distinguish young stem cells from aged stem cells. Here we developed ChromAgeNet, an interpretable deep learning framework trained on 3D DAPI confocal images of HSC nuclei to distinguish between young and aged stem cells.

ChromAgeNet assigns a “youthful score”, representing the likelihood that a nucleus belongs to a young HSC, outperforming a baseline model based on chromatin-extracted features (AUROC 0.77± 0.03 vs. 0.73± 0.04). Our workflow prioritizes biological interpretability and robustness by leveraging the full 3D chromatin architecture, which is an established but underexplored feature in ageing phenotyping. To support generalization, we implemented data augmentations and cross-validation with nucleus-level partitioning. The interpolation along the Z-dimension offered an additional means to synthetically generate 2D slides, preserving realistic distributions of DAPI intensity. Rather than predicting on single 2D slices, we aggregated predictions across Z-stacks, which enhanced performance. To improve reliability of output probabilities, we applied post-hoc beta calibration, aligning youthful scores with true class likelihoods without compromising discriminative power. This work constitutes the first application of deep learning for ageing prediction using 3D microscopy images of HSC nuclei, representing a proof of concept for a scalable and cost-effective platform for high-throughput phenotypic screening of HSC rejuvenating compounds.

Our results revealed substantial phenotypic heterogeneity among HSCs, with not all stem cells from biologically young or aged BM exhibiting the same chromatin characteristics. Individual nuclei from the same age group displayed distinct stages of chromatin architectural alteration or deterioration over time, aligning with previous observations of epigenomic heterogeneity in stem cell ageing [4, 61]. Noteworthily, performance metrics exhibited greater standard deviation across folds for aged HSCs than for young HSCs, reflecting this anticipated increase in phenotypic variability with ageing. In general, we observed differences in classification metrics across folds in both DL and ML approaches, supporting the notion that AI models trained with biological data, which is characterized by inherent variability, should not rely on a single training-validation split, which may introduce biases for model selection. Notably, chromatin-extracted features from 3D nuclear volumes were also effective in discriminating between young and aged HSCs while offering interpretable insights into the ageing-associated archetypes.

SHAP analyses revealed several top-ranked features, mostly associated with textural patterns and particularly from Wavelet and Laplacian-of-Gaussian (LoG) filtered images, that were elevated in aged HSCs and associated with increased entropy, and was consistent with increased chromatin disorganization and loss of epigenomic integrity in the aged nucleus. These findings align with reports of elevated chromatin entropy in aged LADs [31],and an overall loss of chromatin integrity and increased accessibility in aged HSCs [62, 63]. Presumably, the accumulation of stochastic variation in any data modality, which increases entropy readouts, can drive measurements of biological age [64]. Although age did not emerge as a dominant axis of variation in our full feature space, contrary to observations in previous studies in other organs [41], SHAP identified features which contributed to age class separation in UMAP embeddings. While these features exhibited lower predictive power than CNN-based representations, they highlight interpretable and meaningful patterns of ageing-related chromatin remodeling.

To interpret ChromAgeNet’s decisions, we applied XAI methods to identify image regions driving the CNN’s predictions. Among the multiple techniques benchmarked, occlusion sensitivity consistently yielded the most reliable saliency maps. In young HSCs, model attention concentrated along the nuclear envelope, nucleoli, and invagi-nations, while in aged HSCs, attention was more dispersed, often pointing to large DAPI-dense regions and disrupted NE-associated DAPI signals, consistent with LAD deterioration and altered DAPI-Intense Regions [34]. In both age groups, heterochro-matin around nucleoli was highlighted. Interestingly, nucleolar size has been identified as a cellular hallmark of longevity [65, 66]. XAI also revealed fine-textural patterns within the nuclear matrix and higher entropy in attribution maps from aged nuclei, suggesting more diffuse patterns of model attention.

The application of ChromAgeNet to drug-treated aged HSCs demonstrated its potential as a phenotypic readout for rejuvenation drug screening. Rho GTPase pathway inhibitors modestly increased youthful scores, while chromatin-modifying compounds (UNC0646, IOX1) shifted prediction scores closer to those of young HSCs. These results suggest that epigenetic drugs are affecting nuclear features, and might consequently be explored for potential applications to improve the function and regen-erative potential of aged HSCs. While further functional validation is still needed, the relatively small parametric size of ChromAgeNet (approximately 350,000 parame-ters) makes it practical for extending it into a scalable and interpretable image-based platform for screening rejuvenating compounds with single-cell resolution.

Despite the potential of ChromAgeNet for predicting HSC ageing, there are lim-itations to address. First, technical variability in single-cell imaging, stemming from acquisition settings, sample handling, and other experimental parameters, can con-found true biological alterations. Although deep learning models can theoretically learn to ignore the effects of these biases given sufficient data, acquiring large-scale HSC image data is laborious and resource intensive. Our dataset comprises images acquired using two different microscopes and three independent operators. We applied a stan-dardized preprocessing pipeline and data augmentations to enhance model robustness to batch effects, which we later verified by the absence of significant associations between ChromAgeNet estimates and technical parameters **(Supp. Fig. 8**). Nonethe-less, some residual variability may still influence model decisions. Wider application of our model will likely require fine-tuning on independent datasets followed by careful assessment in new experimental contexts. Second, ground truth labels were assigned based on the chronological age of the mouse as a surrogate for biological age. Due to the heterogeneity within aged HSCs discussed previously, this labeling approach introduces learning noise and hinders model convergence at the single-nucleus level. Finally, size-related estimates from DAPI may be biased and could be improved by the use of a more specific nuclear envelope marker, although DAPI borders typically match lamin markers with high confidence [67].

While the exclusive use of DAPI, which is an easily accessible dye, inexpensive and readily compatible with different imaging protocols, represents an advantage of ChromAgeNet, future work could incorporate additional layers of molecular information, for example by staining RNA Pol II and histone post-translational modifications (e.g. H3K9 methylation), which have been shown to refine nuclear phenotyping and improve classification accuracy [41, 47]. Another promising direction is the adoption of 3D CNNs and interpretable methods like 3D GradCAM, although adding one extra dimension and complexity to the learning task demands larger datasets for model training. Larger datasets could also permit the exploration of Visual Transformers to increase prediction performance, although CNNs may remain advantageous for chromatin-based tasks because of their stronger texture bias [68]. Lastly, as single-cell multimodal omics gain traction [69], integrating imaging-derived features into multiomics pipelines represents a complementary and exciting research direction for studying the heterogeneity of cellular ageing.

## 4 Conclusions

We demonstrate that alterations in the nuclear architecture over time can be exploited using a combination of microscopy and ML to construct age prediction models. Our study constitutes a novel use of deep learning algorithms to learn ageing-associated signatures of chromatin changes in DAPI-stained images of single HSC nuclei, as a robust alternative to traditional DNA methylation clocks and other biomarkers of ageing. Despite the intrinsic heterogeneity of ageing and the lack of clear visible differences in DAPI images to the human eye, ChromAgeNet can discriminate young and aged HSCs with confident accuracy. In addition, our XAI pipeline offers causal interpretations of feature model importance at the image-level. Given the absence of HSCs imaging data in public datasets and the labor-intense process of imaging HSCs, we will release our curated 3D HSC microscopy dataset along with ChromAgeNet, offering a valuable open resource to develop and validate computational tools for HSC ageing research, which are currently lacking. The affordability of DAPI and light parametric size of our model make our approach practical for high-throughput microscopy workflows, which could accelerate the discovery of HSC rejuvenating drugs.

## 5 Methods

### Mice

All mice were housed under specific pathogen-free conditions in the animal barrier facility at the Biomedical Research Institute of Bellvitge (IDIBELL) and maintained in accordance with the European Convention for the Protection of Vertebrate Animals used for Experimental and Other Scientific Purposes (ETS 123). All mice were C57BL/6 mice obtained from the internal division stock, where young mice were defined as 10–16 weeks of age, and aged mice as at least 80 weeks old. In the internal division stock, C57BL/6 mice were originally derived from animals sourced from The Jackson Laboratory. Young mice were randomized by sex while aged mice were females. Mice were housed in groups of up to four per cage in Macrolon Type II (long) cages, with bedding and paper nesting material provided. Animals had ad libitum access to food (V1124-3, ssniff®) and water, and were kept under a 12-hour light/dark cycle throughout the experiment.

### HSC extraction and isolation

For obtaining murine HSCs, long bones were harvested and the bone marrow was flushed with a syringe directly into sterile medium. Mononuclear cells were isolated by low-density gradient centrifugation using Histopaque 1083 (Sigma) and stained with a cocktail of biotinylated lineage antibodies. The lineage cocktail included anti-mouse rat antibodies from eBioscience: anti-CD11b (clone M1/70), anti-B220 (clone RA3-6B2), anti-CD5 (clone 53-7.3) anti-Gr-1 (clone RB6-8C5), anti-Ter119 and anti-CD8a (clone 53-6.7). Lineage-positive cells were depleted using magnetic separation with Dynabeads (Invitrogen). Lineage-negative cells were then stained with antibodies against Sca-1 (clone D7), c-Kit (clone 2B8), CD34 (clone RAM34), Flk-2 (clone A2F10), and streptavidin, all from eBioscience. HSCs were gated as Lineage-, c-kit+, sca-1+, CD34-/low, and Flk2-. A representative flow cytometry gating is shown in **Supp. Fig. 7**. HSC were sorted using either a MoFlo-XDP High-Speed Cell Sorter or a CytoFLEX SRT Cell Sorter (both from Beckman Coulter).

### HSC culturing, treatment and imaging

Freshly sorted murine HSCs were seeded onto fibronectin-coated glass coverslips and incubated for 12–16 hours in HBSS supplemented with 10% fetal bovine serum (FBS) and 1% penicillin–streptomycin (P/S). Where indicated, cells were treated with 100 *µ*M Rhosin (RhoAi) [34, 70], 5*µ*M CASIN [33], 50*µ*M IOX1 (8-hydroxyquinoline-5-carboxylic acid), which mimics 2-OG (*α*-KG) and blocks the catalytic activity of lysine demethylases (from Tocris Biotechne), 0.25*µ*M UNC0646 (Sigma), a well-described selective G9a/GLP methyltransferase inhibitor [71] or left untreated. Cells were cultured in growth factor-free medium at 37°C in a hypoxic incubator (5% CO2, 3% O2) and fixed with BD Cytofix fixation buffer (BD Biosciences) following incubation.

Fixed cells were gently washed with PBS and permeabilized with 0.2% Triton X-100 (Sigma) in PBS for 20 minutes. Nuclear staining was performed using DAPI (4;6-diamidino-2-phenylindole) 1*µ*g/*µ*l (Thermo) diluted 1:500 in PBS for 10 minutes, followed by two PBS washes. DAPI is a photostable fluorescent DNA dye and its fluorescence intensity has been used in several applications to quantify DNA amount and chromatin condensation in intact nuclei [72]. Coverslips were mounted using ProLong Gold Antifade reagent without DAPI (Invitrogen, Molecular Probes).

Fluorescence images were acquired with a Zeiss LSM980 or a Zeiss LSM880 confocal microscope with a ×63 objective. Z-stacks were collected by automated scanning along the z-axis, with optical sections acquired every 0.2–0.4 *µ*m. To minimize technical variability, samples were rigorously processed (sorted, stained, and imaged) in parallel within the same experimental batch. Confocal imaging parameters, including laser power, Z-stack interval, and gain settings, were kept constant across all experimental replicates to ensure reproducibility.

### Image preprocessing and segmentation

A total of 1,894 DAPI-stained 3D confocal microscopy images were acquired and exported in Carl Zeiss CZI format, with dimensions of 1,024 × 1,024 pixels in X and Y and a variable number of Z-slices depending on nuclear height, with an average of 29.147 for young HSC and 27.868 for aged HSCs (**Table 1**).

As an initial quality control step, we removed 46 outlier images (∼2% of the dataset) due to overexposure (total pixel intensity *>* 4 · 10^8^) or excessive noise (sigma-estimated noise *>* 8), resulting in a final dataset of 1,848 images, with equivalent distributions among the training classes of young and aged HSCs **(Supp. Fig. 8a,b**).

The remaining images were denoised using Chambolle’s total variation method [73], which outperformed wavelet and bilateral filtering methods in a small validation set, based on the reduction of sigma noise estimates and visual inspection of subnuclear intensity patterns.

To ensure consistent voxel dimensions across samples, we resized each 3D image stack to isotropic resolution (0.1 *µ*m and 0.05 *µ*m per pixel in each X, Y and Z dimension) using interpolation with anti-aliasing, which applies Gaussian smoothing to improve signal-to-noise ratio. A binary nuclear mask was then generated for each image using Otsu thresholding with hysteresis [74], preceded by Gaussian smoothing. Masks were post-processed using hole filling and binary closing operations to correct for segmentation artifacts.

Each intensity image and its corresponding nucleus mask were centered by trimming background regions and applying symmetric padding to approximately match the largest image size across all samples, preserving actual nucleus size and scale. This size is 128 × 128 pixels in XY for 0.1 µm resolution and 224 × 224 pixels in XY for 0.05 *µ*m resolution. Pixel intensities outside the nuclear mask were zeroed out. Final pre-processed 3D arrays were saved in compressed NumPy format (.npz) for efficient data storage and retrieval. These matrices are directly used for manual feature extraction. To generate inputs for ChromAgeNet training, we extracted 2D Z slices from each 3D stack where the binary nuclear mask covered more than 50% of the image pixels. This excluded peripheral slices at the nuclear top and bottom poles, which are uninformative and do not represent the class distribution appropriately. These slices were saved as 8-bit PNG files, an information-preserving format that retains original pixel intensity values.

### Handcrafted chromatin feature extraction

We extract 1,037 handcrafted features related to spatial DAPI chromatin signals from the 3D nucleus images resized to 0.1*µm*/pixel, following definitions established by the Imaging Biomarker Standardization Initiative (IBSI) [75], an initiative to standardize radiomic definitions and enhance reproducibility in these analyses. We define this collection of image-based manually extracted features as *chromatin-extracted features* throughout the text. A schematic graph of our feature extraction and analysis pipeline can be seen in **Supp. Fig. 2a**. Prior to extraction, we applied a set of image filters **(Supp. Fig. 2b**) to enhance spatial intensity features and improve robustness to noise, including Wavelet [76], Laplacian of Gaussian (LoG) filter [77], Gradient and 3D Local Binary Patterns (LBP) [78], all with default parameters.

We set the sigma parameter of the LoG extraction to *σ* = 0.5 and *σ* = 1.0 to capture finer texture structures, as higher sigma values did not produce informative images in our data. Similarly, we also keep only lower pass bands in the first dimension of the Wavelet filter.

Chromatin-extracted features were extracted from the intensity image using the binary nuclear mask, which defined the 3D spatial extent of the nucleus. Image inten-sities within the mask were standardized to zero mean and unit variance to account for inter-sample variability. The extracted features were grouped into three main categories: **morphological**, **intensity-based (first-order)**, and **textural** features. Morphological features included both 2D and 3D shape descriptors that capture geo-metric properties of the nucleus. Intensity-based features measured first-order statistics such as mean, variance, and skewness of pixel intensities within the nuclear mask. Tex-tural features were derived from five established families: Gray-Level Co-occurrence Matrix (GLCM) [79], Gray-Level Difference Matrix (GLDM) [80], Gray-Level Run Length Matrix (GLRLM) [81], Gray-Level Size Zone Matrix (GLSZM) [37], and Neigh-borhood Gray-Tone Difference Matrix (NGTDM) [82]. These matrices quantify spatial relationships, intensity variation, and local homogeneity across the nuclear volume. Full descriptions of all feature categories and how they are computed are available in the PyRadiomics documentation [83].

### Quality Control with chromatin-extracted features

Chromatin-extracted features from young and aged HSC images were standardized using Z-score normalization to have mean=0 and std=1. We identified and removed 134 non-informative features—primarily from LBP3D and Gradient filters—that exhibited constant values across all samples. Furthermore, we found an expected feature redundancy in our radiomics set due to multicolinearity, leading to a notable amount of highly-correlated variables. We apply a filtering to discard those features that exhibit higher pairwise absolute Pearson correlations higher than 0.85 **(Supp. Fig. 2c, Supp. Data Table 1**). The final dataset consisted of 70 numerical features for downstream analysis. The cleaned correlation matrix is shown as a heatmap in **Supp. Fig. 2c**. This feature set was then used to filter bad quality images in a sec-ond quality control step, where the nuclei exhibiting an unusually small (*<* 80*µm*^3^) or large mesh volume (*>* 400*µm*^3^) were filtered out, thus removing 9 outliers. We explored the dataset variability of young and aged nuclei conditions after dimen-sionality reduction with Uniform Manifold Approximation and Projection (UMAP) **(Supp. Fig. 9a,b**), to assess that no technical variable is affecting the overall data variance. We identified a UMAP region with images that exhibited increased values for technical variables such as raw image mean intensity and sigma noise **(Supp. Fig. 9c,d**). We characterized this population of nuclei through K-Means clustering and discarded them from subsequent analyses **(Supp. Fig. 9e**), similarly to how it has been done in single-cell studies [84]. We decided to use this approach instead of regressing out confounding variables to not distort original feature values. Re-running the UMAP with the filtered data revealed no sign of aggregation of these technical signals **(Supp. Fig. 9f-h**), and the spread of technical variables and batches was more homogeneous in the latent space, successfully minimizing the effect of these technical biases over the total dataset variability.

### ML analyses of chromatin-extracted features

To assess whether chromatin-extracted features could distinguish young from aged HSCs, we benchmarked four ML classifiers — Logistic Regression (LR), Random Forests (RF), XGBoost, and Calibrated XGBoost (Cal. XGBoost) — to evaluate the classification performance of our chromatin feature set. Each model was tested in combination with different feature selection strategies: correlation-based selection using ANOVA F-test, Recursive Feature Elimination (RFE), and model-based feature importance. To correct the overconfident probability estimates produced by XGBoost, we applied isotonic calibration [85] within an inner 5-fold cross-validation loop. Model performance was assessed using an outer 5-fold cross-validation stratified by class labels, reporting accuracy, precision, recall, AUC, and F1 score for each configuration.

At the start of each cross-validation loop, feature standardization was performed on the training data and applied to the corresponding validation fold. Feature selection was also restricted to the training set to avoid information leakage and subsequently applied to both training and validation splits. Hyperparameters such as the number of estimators and maximum depth (for RF and XGBoost) were optimized using a randomized grid search over 100 iterations, evaluated by 4-fold inner cross-validation **(Supp. Table 7**). For RFE, the optimal number of features was determined by iteratively training models with feature subsets ranging from 10 to 50 and selecting the configuration yielding the best performance **(Supp. Fig. 3d**). All experiments were conducted using a fixed random seed to ensure reproducibility and enable consistent, fair comparison across models and feature selection strategies.

Feature importance in the best-performing model was assessed using SHAP (SHapley Additive exPlanations) values computed with the TreeExplainer() function [52]. SHAP is a model-agnostic XAI method that assigns an importance value to each feature in a model by comparing a model’s output with and without an individual feature. We report SHAP values for the top-ranked features in each cross-validation fold, alongside AUROC curves, precision-recall curves, and confusion matrices. To generate the final UMAP embedding, we identified the intersection of the most important SHAP features across all folds, resulting in a feature set of 15 common features.

### Data partitions for learning the network

All samples from young (n = 551) and aged (n = 552) classes were shuffled and partitioned into five folds for cross-validation (CV). Aged HSC images in DMSO medium (n = 126) were evenly distributed across folds as part of the aged class. In each iteration, four folds were used for adjusting the model’s weights and the remaining fold for validation. Splits were performed at the 3D nucleus level to ensure that all 2D slices from a given nucleus remained within the same fold, preventing data leakage. Each fold contained approximately 16,000 2D images (∼240 nuclei) and was stratified by age class to maintain a consistent 45:55 ratio of young to aged cells. For supervised learning, images with HSCs nuclei from young mice were labeled as y=1 and those from aged mice as y=0.

### ChromAgeNet architecture

ChromAgeNet network architecture is a reduced adaptation of the Xception (Extreme Inception) network, which leverages innovative CNN modules such as depthwise separable convolutions and residual connections with a linear bottleneck structure. Our network architecture is the following. ChromAgeNet input layer, with dimensions 128 × 128 × 1, is followed by a 2D convolution with stride S=2, followed by Batch Normalization. The output is passed through a sequence of repeated blocks consisting of depthwise separable convolutions with 10% spatial dropout, each followed by 2D average pooling to downsample the spatial dimensions. Batch Normalization is applied after each convolutional layer. We alternate between swish and leaky ReLU activation functions after each convolution throughout the network. Convolutional kernels remained constant through the network, with sizes (3 × 3) and “Same” padding. The number of channels increases progressively across the network as [32, 64, 64, 128, 256, 256]. After the final convolutional block, global spatial information is aggregated via an AveragePooling2D layer into a feature vector. This is followed by three fully connected layers of sizes [256, 64, 16] with ReLU activation, and a final output unit with a sigmoid function that returns the probabilistic score. This combination of hyperparameters resulted in 347,393 trainable parameters and 2,624 non-trainable parameters.

### ChromAgeNet training and inference

ChromAgeNet is trained to output a probability reflecting how likely a given 2D image corresponds to a young nucleus, and the score is aggregated for all images belonging to the same nucleus to obtain a final nucleus-level score. Individual images were loaded in batches of 64 and processed with on-the-fly data augmentation applied exclusively to the training set. Each dataset partition was shuffled, batched, cached, and preloaded using prefetching to enable efficient in-memory operations and minimize input/output latency during training. Augmentations included random horizontal and vertical flips, random rotations (with background filling), and contrast adjustments by a small factor of ±0.05. These transformations help reduce the influence of technical variation and batch effects in image acquisition by increasing sample variability. Additionally, the isotropic interpolation applied during preprocessing effectively acts as a form of data augmentation by synthesizing intermediate 2D slices from 3D volumes.

Model weights were optimized using the AdamW optimizer with a learning rate of 0.0001. We applied an Exponentially Moving Average (EMA) to stabilize training and improve generalization. We use Binary Cross-Entropy as the loss function with 0.1 label smoothing to learn the network weights and apply Just-In-Time (JIT) compilation to accelerate model training and inference. We set the same initial weights at the beginning of each cross-validation fold to discard model differences due to random weight initialization. In all experiments, validation loss did not converge consistently across epochs, and we therefore applied Early Stopping to prevent overfitting.

Because we aim to maximize the separation of the two classes to calibrate a probabilistic range, we fixed the Area Under the Receiver Operating Characteristic (AUROC) as the primary optimization metric, as it reflects the model’s ability to separate these classes across varying decision thresholds. The optimal classification threshold was determined by maximizing Youden’s J statistic (computed as J = TPR – FPR) from the ROC curve of each cross-validation fold, to optimize the trade-off between sensitivity and specificity. Additional metrics including accuracy, precision, recall, AUC, and F1 score are reported using the mean, standard deviation, and 95% confidence interval across all folds.

To generate the prediction of the model at the nucleus level, we aggregate image-level predictions from Z-slices belonging to the same nucleus using soft-voting. Soft voting computes the average probability across all 2D image-level predictions for a given nucleus:

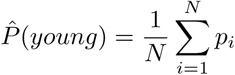

Where *P̂*(*young*) is the ChromAgeNet probability for class *y* = 1 at the 3D nucleus level, *N* is the total number of 2D images per nucleus and *p_i_* is the predicted probability for class *y* = 1 for the *i*-th 2D image. The final nucleus class prediction is obtained by selecting a class threshold *t*:

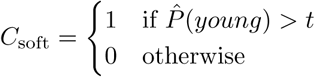

We made sure ChromAgeNet is not driven by major technical sources of variability in our dataset. ChromAgeNet scores were consistent for aged and young HSCs across microscopist and number of dyes performed on the sample **(Supp. Fig. 8c,d**). Furthermore it showed no strong correlations to technical variables of the image meta-data such as mean pixel intensity, total pixel intensity, estimated noise or number of Z stacks **(Supp. Fig. 8e-h**).

### Model Calibration

The main goal of ChromAgeNet training is to calibrate a probability range between the learned distributions of aged and young HSCs, where properly calibrated probabilities are crucial.

ChromAgeNet classifies HSCs as young or aged based on chromatin patterns, with well-calibrated probabilities being essential for downstream analyses such as rejuvenation scoring for drug discovery applications. We calibrated ChromAgeNet’s predictions using the validation fold in each cross-validation iteration to ensure that predicted probabilities reflect true class likelihoods. To quantify calibration quality, we used reliability curves, the Expected Calibration Error (ECE), that measures solely the model calibration, and the Brier Score (BS), a metric that captures both the accuracy and calibration of probabilistic predictions. Lower ECE and BS indicate better-calibrated probabilities that are closer to the actual class distributions. We used Scikit-learn’s implementation for the Brier Score, and computed ECE as follows:

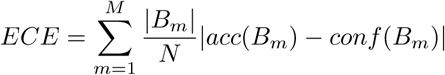

Where M is the number of bins, N the total number of samples, *B_m_* is the set of samples where the predicted probability falls into the *m*-th bin, and |*B_m_*| is the number of samples in bin *B_m_*. *acc*(*B_m_*) is the empirical accuracy in bin *B_m_*, while *conf* (*B_m_*) refers to the mean predicted probability in bin *B_m_*.

This calibration strategy learns a post-hoc mapping over the model scores to align them to real data frequencies. We decided to apply Beta Calibration [51] as its less prone to overfit on small datasets than its non-parametric counterpart, isotonic regression, and has been proven to be an improved method to logistic calibration (Platt Scaling). The final calibrated model was applied subsequently to all the analyses specified in the article for inference.

### Explainability analyses

To interpret the image representations learned by ChromAgeNet, we applied XAI methods to the best-performing cross-validation split. Activation maximization [53] synthesizes inputs to maximize the response of a given convolutional filter, revealing hierarchical patterns learned across network layers. This technique was performed by iteratively optimizing an input image to maximize the activation of a selected convolutional filter using gradient ascent (30 iterations, learning rate = 10), with L2-normalized gradients at each step. For the remaining analyses, we substituted the final sigmoid activation function to a linear function, as generally recommended, as it confers more stable gradient by eliminating non-linearity and operating directly on logits.

To interpret predictions on real data, we used attribution methods such as SHAP [52], which assign importance scores to input pixels based on their contribution to the model’s output. Input data consisted of 1,000 high-confidence correctly predicted images per class (p(young) *>* 0.9 for young HSC and p(young) *<* 0.1 for aged HSC) from the validation set. SHAP values were computed using the Explainer() class from the SHAP Python library, with max_evals=500 and batch_size=50.

For more general attribution benchmarking, we evaluated multiple explainability methods implemented in Xplique [54], using default settings and a batch size of 64. Attribution method comparisons were performed using MuFidelity [55], as implemented in the same library, which evaluates how well the attribution scores correlate with the model’s sensitivity to input perturbations. Among all methods, Occlusion achieved solid high fidelity scores among classes, specially in aged HSCs **(Supp. Table 8**). Occlusion sensitivity [86], a model-agnostic algorithm, measures pixel importance by sliding a small rectangular occluding patch across the image and quantifies its effect on the model prediction. Similarly, Hilbert-Schmidt Independence Criterion (HSIC) attention maps were plotted for comparison, covering a wider area in pixel space [56]. Attribution maps were overlaid on original intensity images, with attention values clipped between the 0.5th and 99.5th percentiles.

To systematically assess attention, Shannon entropy was computed on attribution maps using scikit-image [87] while attribution values by distance to the nuclear border was computed using the distance transform from scipy library. Here, the attribution intensities within each distance value (integer corresponding to number of pixels) were summed and then averaged at the image level.

### Statistical Tests

Statistical significance was performed using two-sided Mann-Whittney U-test and corrected for multiple comparison using Bonferroni method unless specified otherwise.

### Software and versions

ChromAgeNet code was developed using a Python v3.12.7 environment. We took advantage of the AICSImageIO package [88] v4.14.0 with AICSpylibczi v3.2.0 to parse the raw image data and metadata contained in the .CZI files. The image operations were performed with Scikit-Image v0.24.0 [87]. We used Keras [89] to build, train, and validate our convolutional neural network, using TensorFlow v.2.17.0 as the back-end [90]. We employed PyRadiomics v3.0.1 [83] to extract morphological, intensity and texture-related features from 3D nuclear matrices. Statsannotations v0.6.0 [91] was used to report statistical significance. For the AI explanation section, we used the methods of UMAP v0.5.6 [92], SHAP values v0.47.1 [52], and Xplique v1.4.0 [54]. We also used Numpy v1.26.4 [93], Seaborn v0.13.2 [94], Scikit-learn v1.5.2 [95], and Pandas v2.2.3 [96] for numerical operations, plotting, machine learning development, and dataframe management, respectively. Other packages may have been used for minor tasks.

## Supplementary information

- **Supplementary Figures (supplementary figures.pdf):** PDF Document containing the supplementary figures referenced throughout the main text.
- **Supplementary Tables (supplementary tables.pdf):** PDF Document containing the supplementary tables referenced throughout the main text.
- **Supplementary Data 1 (SuppData1.Filtered features.csv):** CSV file containing the values of 70 selected features extracted from the segmented 3D intensity images of individual nuclei.
- **Supplementary Data 2 (SuppData2.SHAP.csv):** CSV file containing the SHAP values for our image-derived feature set for each of the validation folds during cross-validation.

## Declarations

### Ethics approval

All animals were maintained following the European Convention guidelines for the Protection of Vertebrate Animals used for Experimental and other Scientific Purposes (ETS 123). All experiments involving mice were conducted in accordance with the ethical standards set by the Spanish Law for Animal Protection and Welfare Code, having been previously approved in the project AR18008/10399 by IDIBELL’s Ethical Committee for Animal Experimentation (CEEA-IDIBELL) along with approval from the Generalitat de Catalunya.

### Availability of data and materials

The datasets generated and analysed during the current study are available in the Catalan Open Research Area (CORA) repos-itory https://doi.org/10.34810/data2372. This link and the code repository hosting all source code developed for this project will be openly available after publication.

### Author contribution

P.I.P. developed the software presented in this study and wrote the manuscript. E.M., D.B., E.V. performed animal experiments and collected imaging data. P.I.P., M.C.F. and P.P. contributed to the study design of analyses and experiments. M.C.F. and P.P. conceptualized the study, co-supervised the work, revised, and edited the manuscript. All authors have read and agreed to the published version of the manuscript.

## Acknowledgments

We thank the members of the Biomedical Data Science at ISGlobal and the Stem Cell Aging lab at IDIBELL for helpful advice and discussions. We thank ISGlobal SRI Unit for providing cloud computing support and general technical assistance throughout the project, and Sara Montserrat Vázquez for helping setting up the dataset repository. We thank CERCA Program/Generalitat de Catalunya for institutional support.

