## Supplementary material for "Deep learning predicts haematopoietic stem cell ageing from 3D chromatin images": Supp. Fig.

#### Supplementary figure 1

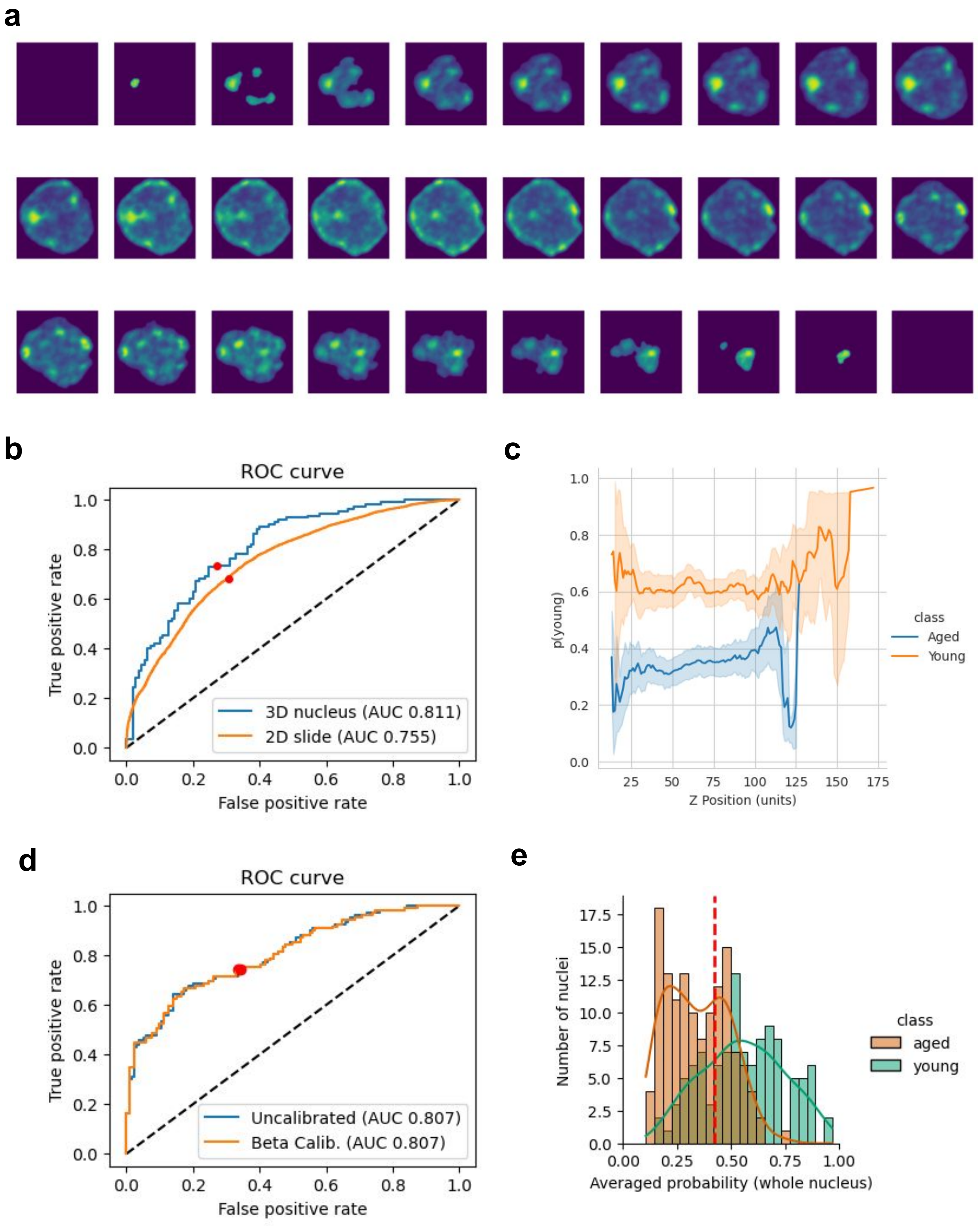

**Supplementary figure 1**

**a.** Series of 2D images in the XY plane depicting several Z-stacks for the same nucleus in ascending order, from bottom to top. **b.** ROC curves, representing the True Positive Rate vs. False Positive Rate on the validation dataset before and after soft-voting for the best performing cross-validation fold. The classification threshold yielding the highest AUC score is shown as a red dot. **c.** Lineplots depicting mean probability scores over each Z-stack position in the 3D nucleus for each class, error bar depicts 95% CI. Distributions of ChromAgeNet scores for the validation dataset after calibration. **d.** ROC curve and AUC for ChromAgeNet performance on validation data for one CV fold before and after calibration. **e.** Distributions of ChromAgeNet scores for the two ground-truth classes in the validation dataset after applying Beta-Calibration on the validation set.

### Supplementary figure 2

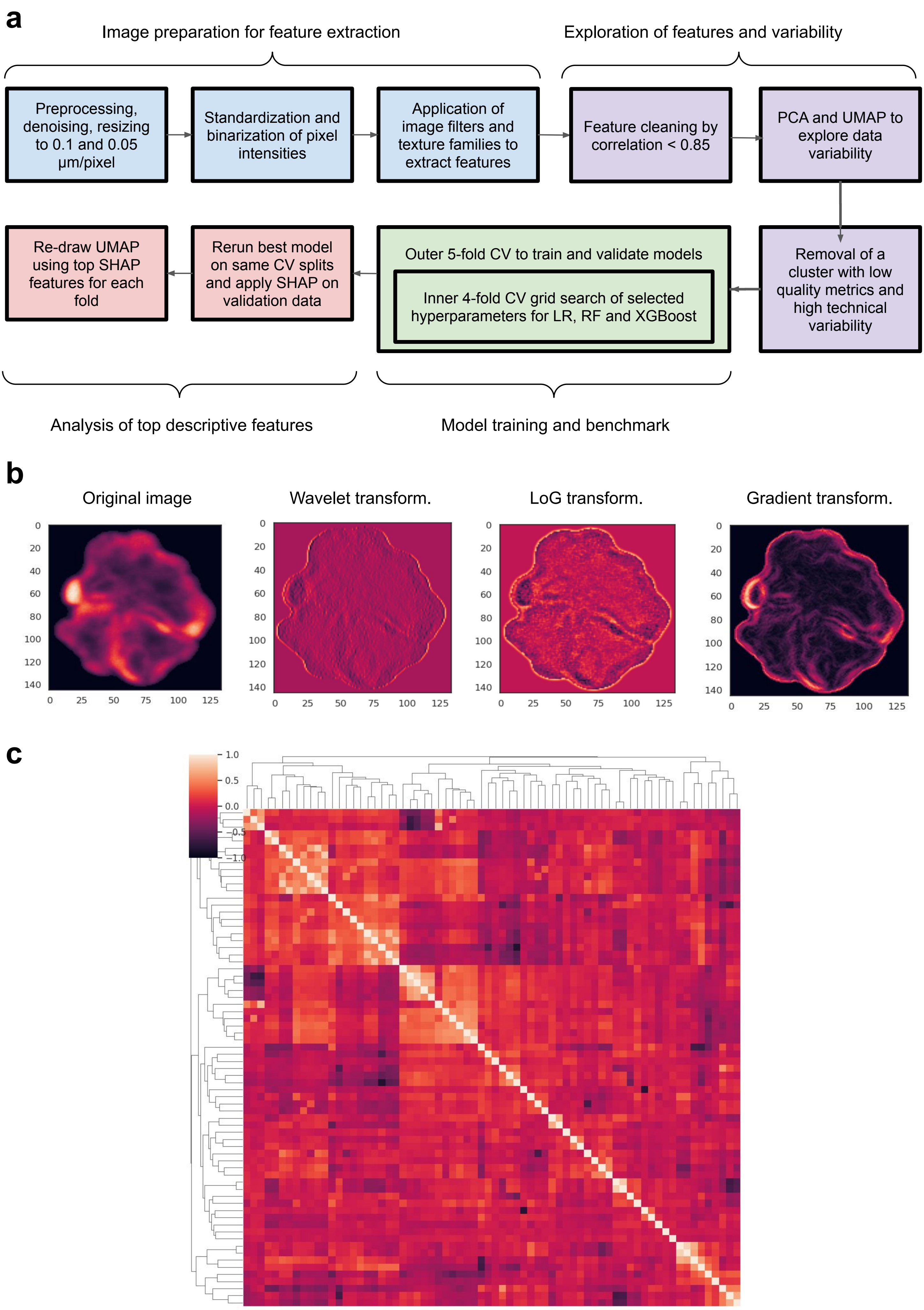

#### Supplementary figure 2

**a.** Schematic representation of our image preprocessing, manual feature extraction and analysis, and ML modelling **b.** Representative images for the product of different filters applied to a HSC nucleus image. **c.** Correlation matrix of the extracted feature set, represented as a heatmap after performing filtering by correlation > 0.85.

### Supplementary figure 3

**a**

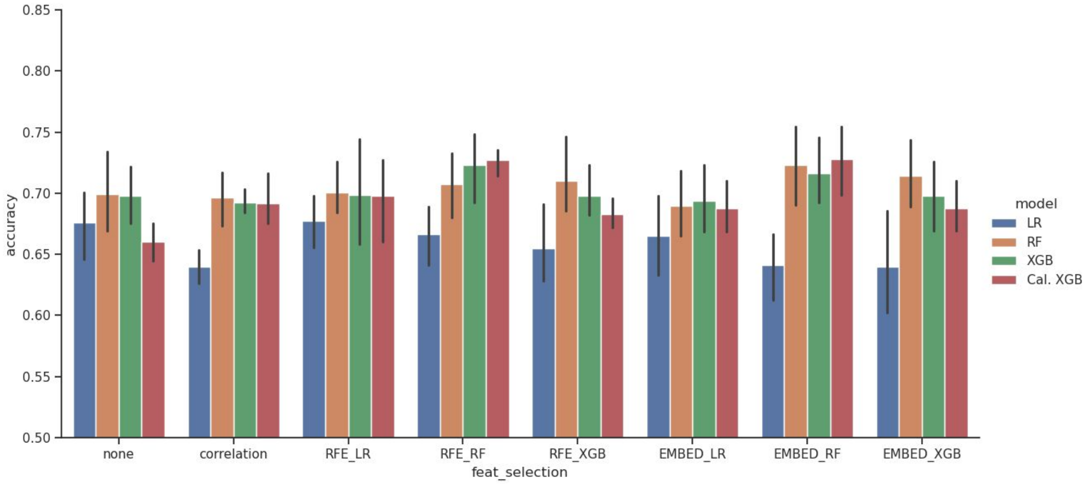


**b**

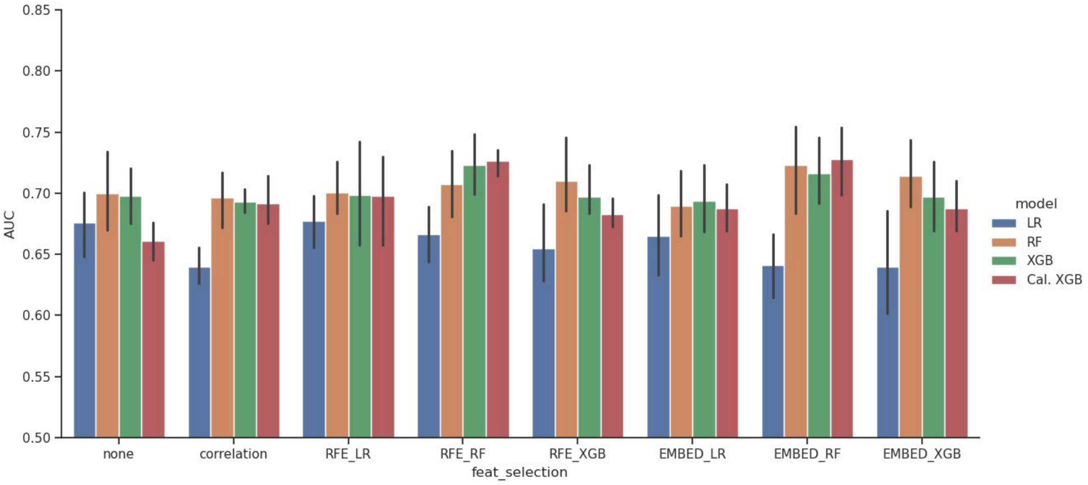


**c**

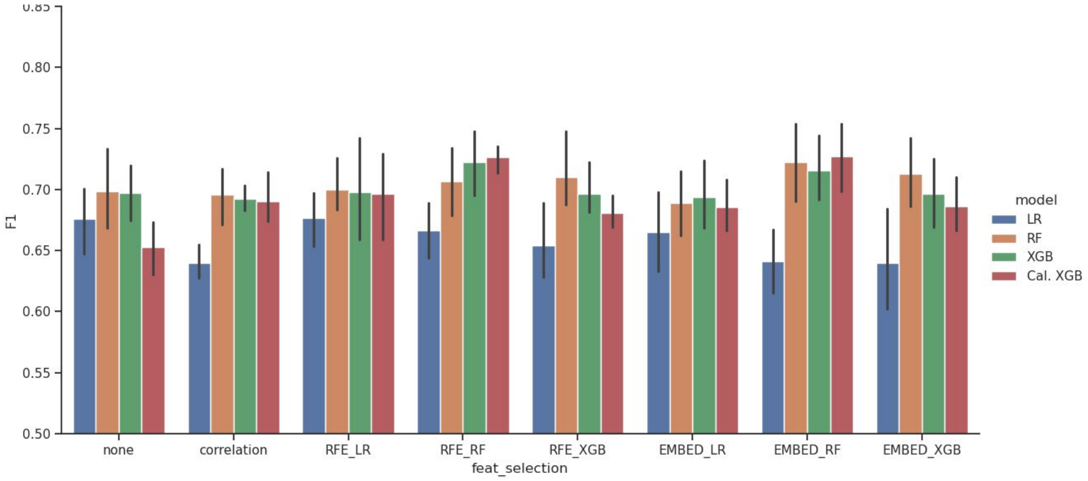


**d**

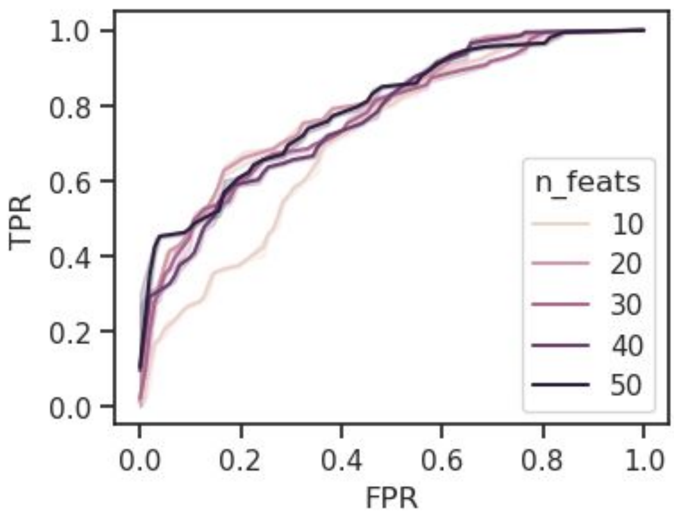

#### Supplementary figure 3

**a-c.** Boxplots representing the comparison of performance metrics from the benchmark among different models and feature selection methods over our radiomics feature set. The metrics are accuracy (top), ROC-AUC (middle) and F1 (bottom). The error bar depicts the standard deviation of each metric among the 5 different cross-validation iterations. **d.** ROC-AUC curve for XGBoost model over different selected number of features during Recursive Feature Elimination (RFE).

### Supplementary figure 4

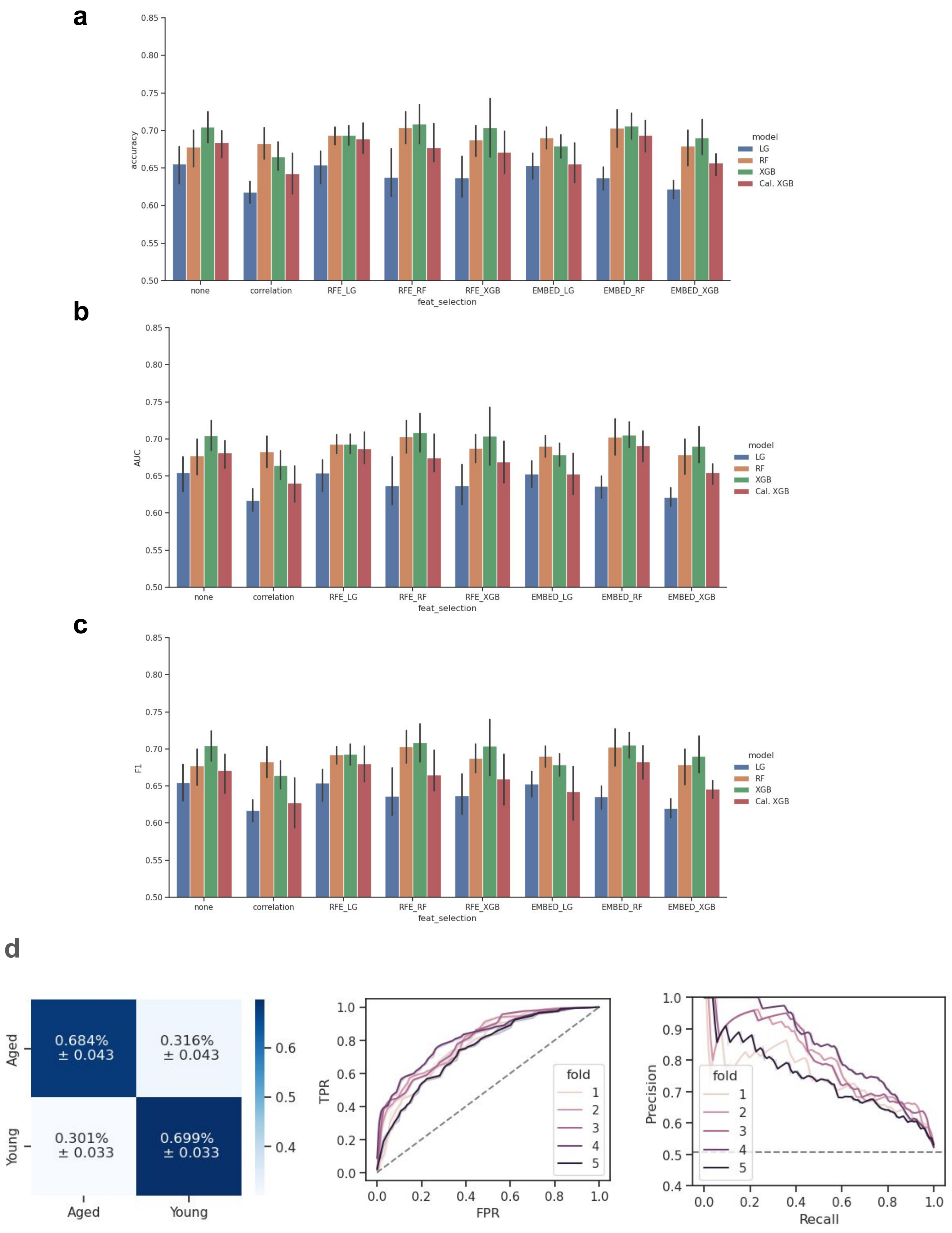

#### Supplementary figure 4

**Model performance on extracted features from binarized images. a-c.** Boxplots representing the comparison of performance metrics from the benchmark among different models and feature selection methods over our handcrafted feature set for images that have been previously binarized for intensity. The metrics are accuracy (top), ROC-AUC (middle) and F1 (bottom). The error bar depicts the standard deviation of each metric among the 5 different cross-validation iterations. **d.** Plots showing the performance of our best model, including the confusion matrix averaged over all folds, and the ROC and Precision-Recall curves for each of the data splits in our cross-validation pipeline.

### Supplementary figure 5

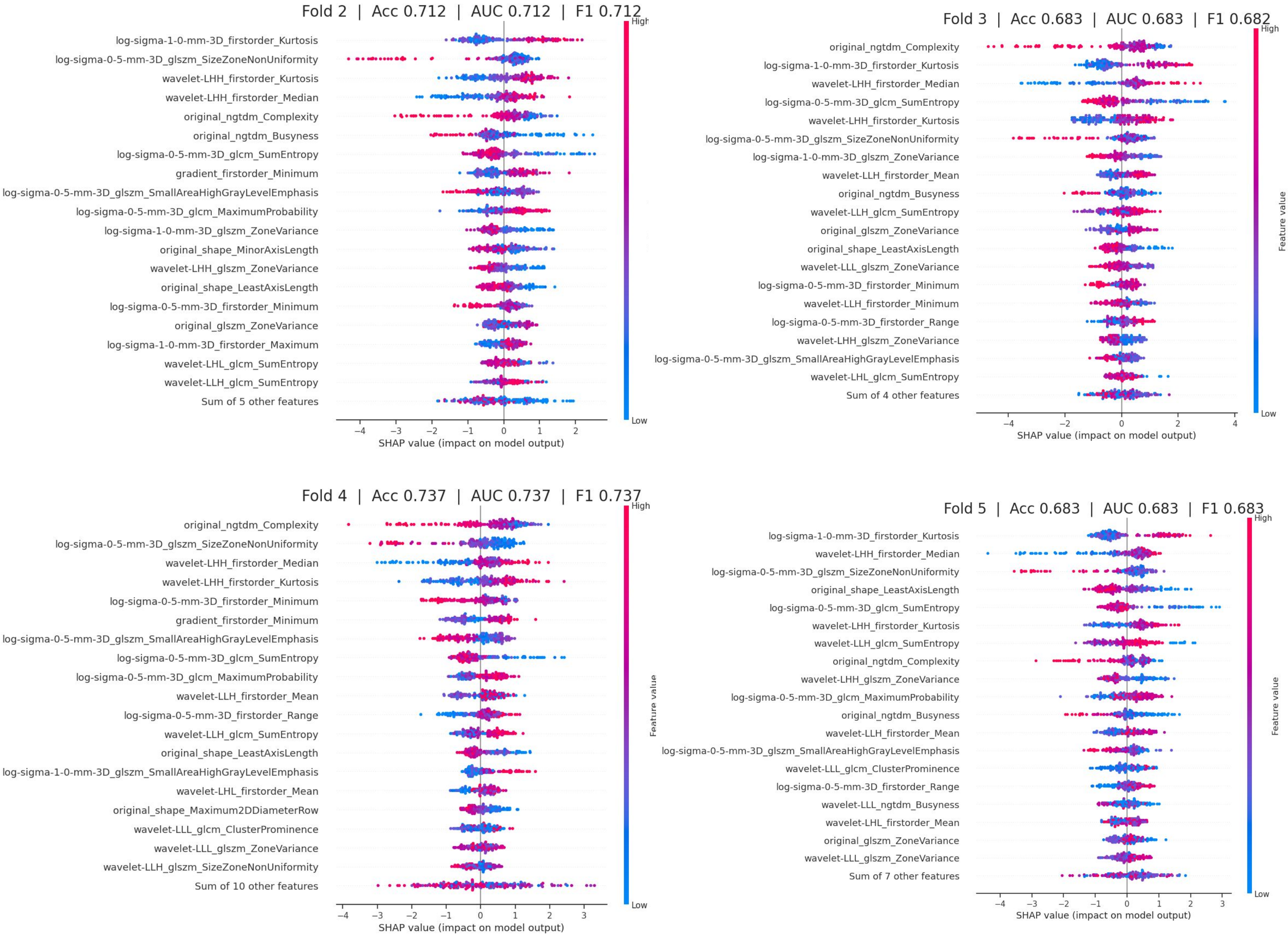

**Supplementary figure 5**

Summary plot of feature importance calculated using SHAP values for cross-validation folds 2 to 5. The X axis represents the impact of each model on the model's output (contribution toward the young HSC class), while the color shows the scaled feature value (higher values are colored as red, while lower values are colored as blue).

**Supplementary figure 6**

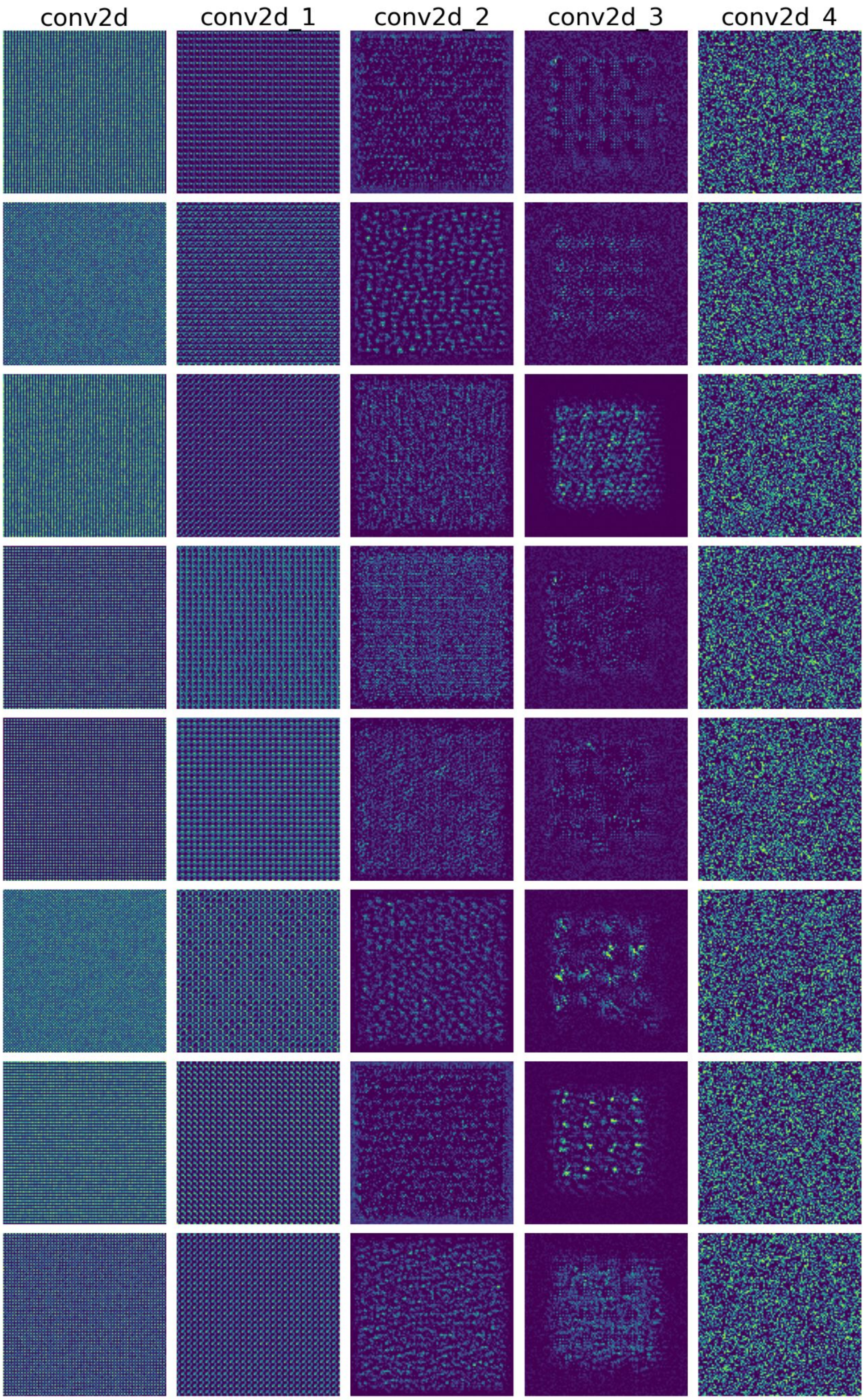

**Supplementary figure 6**

Sequence of Activation Maximization (AM) plots, each column corresponding to a different convolutional layer of our network, and each row to a different convolutional unit. The complexity of the maps learned increase as we move deeper in the network, with an association of reduction of noise. The maps from the 4th do not exhibit intensity near the image border, and demonstrate higher focus to the center of the image where the nucleus is. Interestingly, the 5th layer produced AM plots with a disorganized texture representing chromatin patterns that covers the whole image.

### Supplementary figure 7

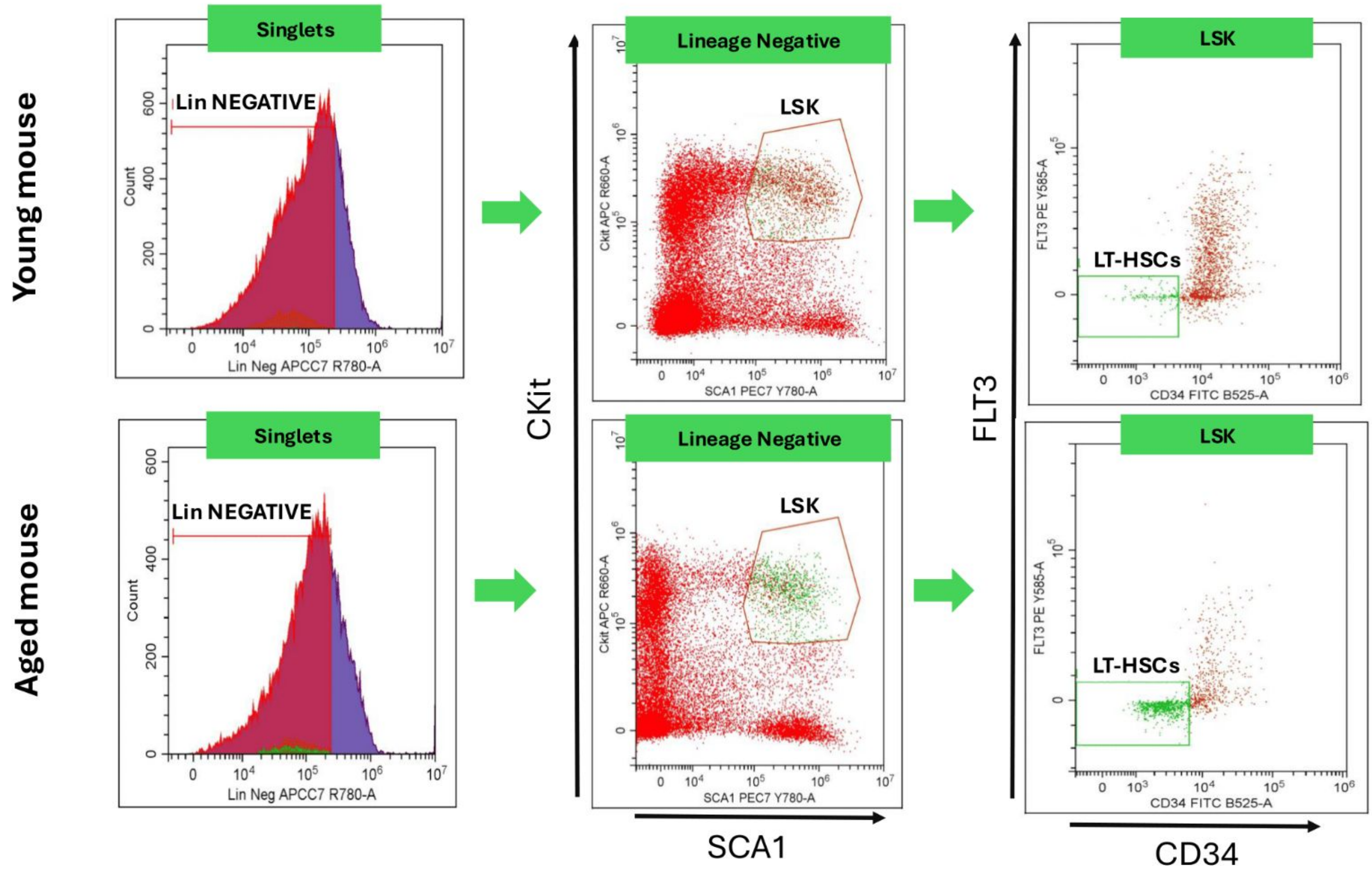


Representative gating strategy for LT-HSCs (Lin- c-kit+ Sca-1+ Flk2- CD34-) sorted from young and aged C57Bl6 mice.

### Supplementary figure 8

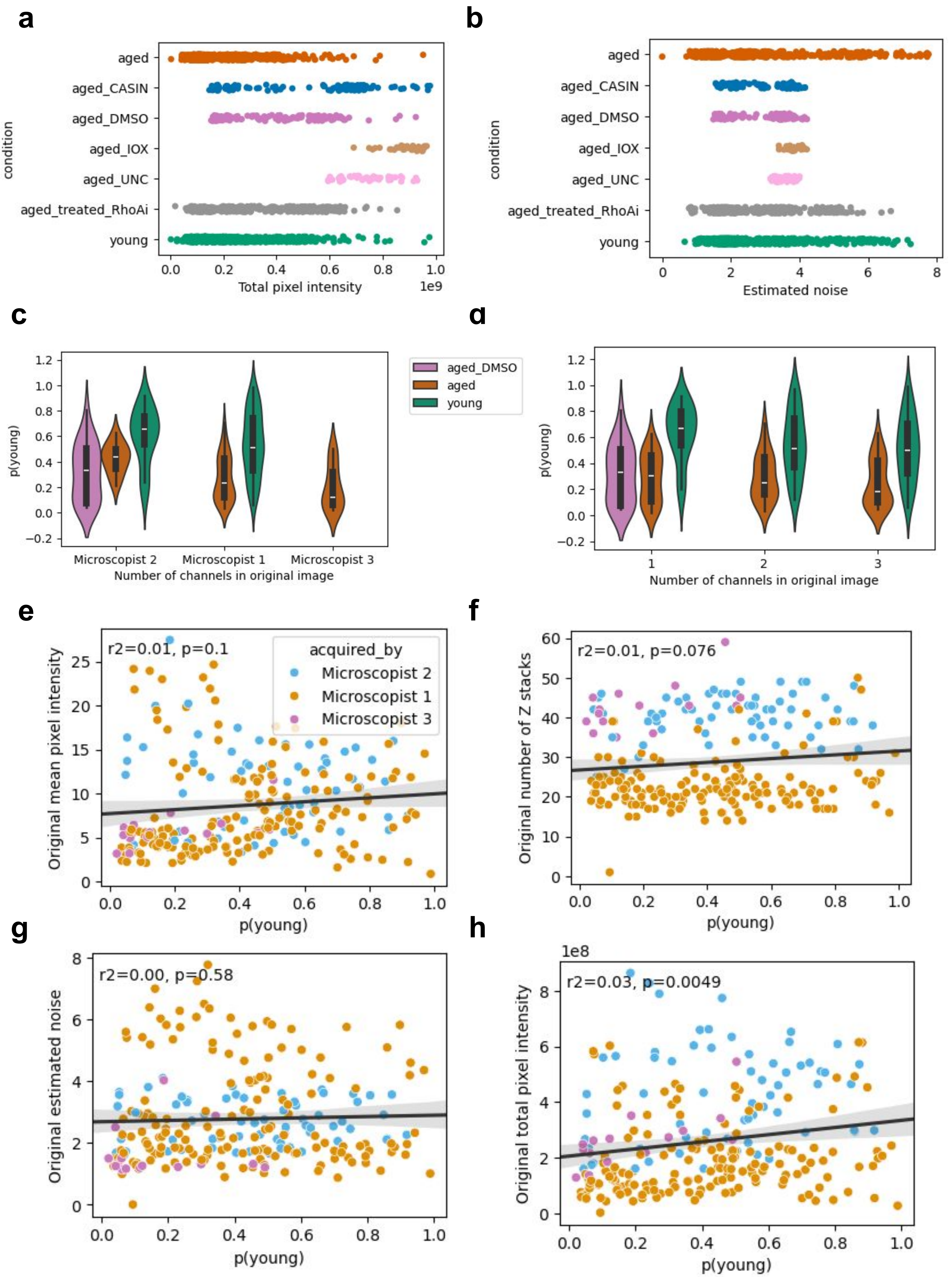

**Supplementary figure 8**

**a-b.** Distribution by biological condition of technical variables in the raw data, with total pixel intensity (**a**) and estimated noise (**b**). **c-d.** Violinplots showing ChromAgeNet scores for validation data classes, segregated across different microscopists (**c**) and fluorochrome dyes applied to the sample (**d**). **e-h.** Scatterplots depicting ChromAgeNet scores in the x axis and numerical metadata variables in the y axis, with each dot representing an individual HSC nucleus. A black regression line shows the relationship among the two, with the strength of the correlation shown as Pearson coefficient and p-value on top of each plot, which indicates the proportion of variance in one variable that can be explained by the other variable. Dots are colored by microscopist.

### Supplementary figure 9

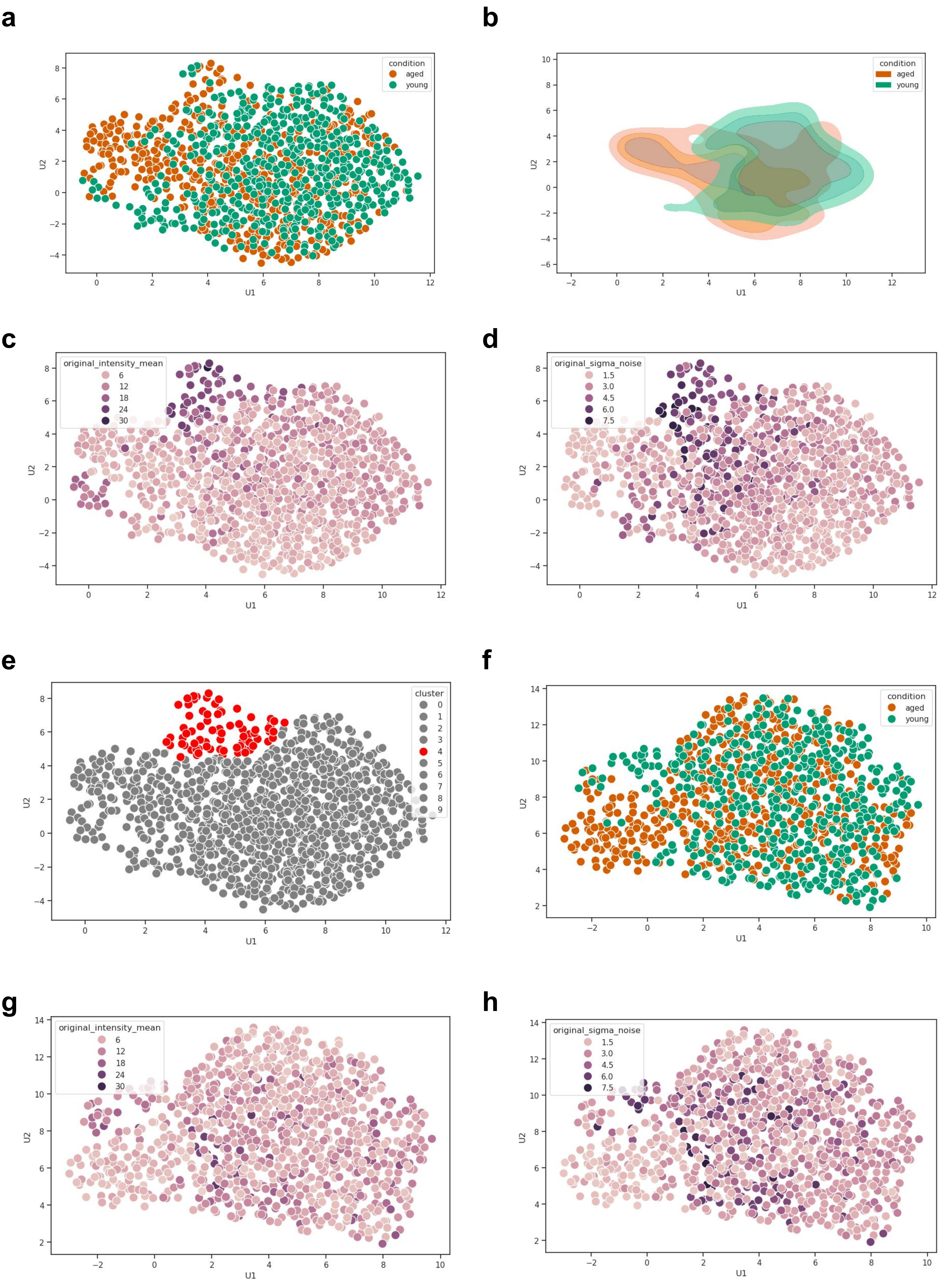

#### Supplementary figure 9

**a, b.** UMAPs computed using all the images, depicting the separation among young and aged HSCs as a scatterplot **(a)** and kernel-estimated densities **(b)** **c, d.** Same UMAP colored by the mean intensity in the original raw images **(c)** and by the estimated noise in the original raw images. **e.** UMAPs computed using all the images, highlighted by the identified cluster with lower QC images using K-Means clustering. **f.** UMAPs computed with the filtered set of images, depicting the separation among young and aged HSCs as a scatterplot **g, h.** UMAPs computed using the filtered set of images, colored by the mean intensity in the original raw images **(g)** and by the estimated noise in the original raw images.
