## Supplementary material for "Deep learning predicts haematopoietic stem cell ageing from 3D chromatin images": Supp. Table

Table 1: Performance Metrics of ChromAgeNet over the 5 cross-validation folds at 2D XY slide level.

| Fold | Accuracy | Precision | Recall | AUC | F1 |
|---|---|---|---|---|---|
| 1 | 0.69 | 0.63 | 0.68 | 0.76 | 0.66 |
| 2 | 0.62 | 0.54 | 0.81 | 0.70 | 0.65 |
| 3 | 0.68 | 0.65 | 0.62 | 0.73 | 0.63 |
| 4 | 0.65 | 0.60 | 0.69 | 0.72 | 0.64 |
| 5 | 0.65 | 0.59 | 0.70 | 0.71 | 0.64 |
| All mean | 0.66 | 0.60 | 0.70 | 0.72 | 0.64 |
| All std | 0.03 | 0.04 | 0.07 | 0.02 | 0.01 |
| All CI | (0.63, 0.68) | (0.57, 0.64) | (0.64, 0.76) | (0.70, 0.74) | (0.64, 0.65) |

Table 2: Performance Metrics of ChromAgeNet over the 5 cross-validation folds at nucleus level using soft-voting.

| Fold | Accuracy | Precision | Recall | AUC | F1 |
|---|---|---|---|---|---|
| 1 | 0.70 | 0.64 | 0.74 | 0.81 | 0.69 |
| 2 | 0.60 | 0.53 | 0.90 | 0.74 | 0.66 |
| 3 | 0.72 | 0.70 | 0.64 | 0.78 | 0.67 |
| 4 | 0.71 | 0.65 | 0.80 | 0.76 | 0.72 |
| 5 | 0.68 | 0.61 | 0.80 | 0.76 | 0.69 |
| All mean | 0.68 | 0.63 | 0.78 | 0.77 | 0.69 |
| All std | 0.05 | 0.06 | 0.09 | 0.03 | 0.02 |
| All CI | (0.64, 0.72) | (0.57, 0.68) | (0.70, 0.86) | (0.75, 0.79) | (0.67, 0.70) |

Table 3: Performance Metrics of Feature Selection Methods for the classification of manually-extracted features. Shown as Average ± Std (CI).

| Method | Accuracy | Precision | Recall | AUC | F1 |
|---|---|---|---|---|---|
| EMBED LR | 0.68 ± 0.03 (0.65, 0.71) | 0.69 ± 0.04 (0.65, 0.72) | 0.69 ± 0.06 (0.64, 0.73) | 0.68 ± 0.03 (0.65, 0.71) | 0.68 ± 0.03 (0.65, 0.71) |
| EMBED RF | 0.70 ± 0.05 (0.66, 0.75) | **0.70 ± 0.06 (0.65, 0.75)** | 0.70 ± 0.05 (0.66, 0.75) | 0.70 ± 0.05 (0.66, 0.75) | 0.70 ± 0.05 (0.66, 0.75) |
| EMBED XGB | 0.69 ± 0.05 (0.65, 0.73) | 0.68 ± 0.05 (0.64, 0.73) | 0.70 ± 0.07 (0.64, 0.76) | 0.69 ± 0.05 (0.65, 0.73) | 0.68 ± 0.05 (0.64, 0.72) |
| RFE LR | 0.69 ± 0.04 (0.66, 0.73) | 0.69 ± 0.04 (0.65, 0.73) | 0.71 ± 0.05 (0.67, 0.75) | 0.69 ± 0.04 (0.66, 0.73) | 0.69 ± 0.04 (0.66, 0.73) |
| RFE RF | **0.71 ± 0.04 (0.67, 0.74)** | **0.70 ± 0.04 (0.67, 0.74)** | **0.72 ± 0.05 (0.67, 0.76)** | **0.71 ± 0.04 (0.67, 0.74)** | **0.71 ± 0.04 (0.67, 0.74)** |
| RFE XGB | 0.686 ± 0.037 (0.65, 0.72) | 0.69 ± 0.04 (0.65, 0.72) | 0.69 ± 0.07 (0.64, 0.75) | 0.69 ± 0.04 (0.65, 0.72) | 0.69 ± 0.04 (0.65, 0.72) |
| Correlation | 0.68 ± 0.03 (0.65, 0.71) | 0.68 ± 0.04 (0.65, 0.72) | 0.68 ± 0.05 (0.64, 0.72) | 0.68 ± 0.03 (0.65, 0.71) | 0.68 ± 0.03 (0.65, 0.71) |
| No selection | 0.68 ± 0.03 (0.65, 0.71) | 0.68 ± 0.04 (0.64, 0.72) | 0.70 ± 0.08 (0.63, 0.77) | 0.68 ± 0.03 (0.65, 0.71) | 0.68 ± 0.04 (0.65, 0.71) |

Table 4: Performance Metrics of shallow ML Models for the classification of manually-extracted features. Shown as Average ± Std (CI).

| Fold | Accuracy | Precision | Recall | AUC | F1 |
|---|---|---|---|---|---|
| LR | 0.66 ± 0.04 (0.63, 0.69) | 0.66 ± 0.04 (0.62, 0.69) | 0.66 ± 0.04 (0.62, 0.70) | 0.66 ± 0.04 (0.63, 0.69) | 0.66 ± 0.04 (0.63, 0.69) |
| RF | **0.71 ± 0.03 (0.68, 0.74)** | 0.70 ± 0.04 (0.67, 0.73) | **0.72 ± 0.05 (0.68, 0.77)** | **0.71 ± 0.03 (0.68, 0.74)** | **0.71 ± 0.03 (0.67, 0.74)** |
| XGB | 0.70 ± 0.03 (0.67, 0.73) | **0.70 ± 0.04 (0.67, 0.74)** | 0.71 ± 0.04 (0.67, 0.74) | 0.70 ± 0.03 (0.67, 0.73) | 0.70 ± 0.03 (0.67, 0.73) |
| Cal. XGB | 0.70 ± 0.03 (0.67, 0.72) | 0.70 ± 0.05 (0.66, 0.74) | 0.70 ± 0.08 (0.64, 0.77) | 0.69 ± 0.03 (0.667, 0.72) | 0.69 ± 0.04 (0.66, 0.72) |

Table 5: Performance metrics across different folds for all shallow ML models for the classification of manually-extracted features. Shown as Average ± Std (CI).

| Fold | Accuracy | Precision | Recall | AUC | F1 |
|---|---|---|---|---|---|
| 1 | **0.73 ± 0.03 (0.70, 0.76)** | **0.73 ± 0.04 (0.69, 0.77)** | **0.74 ± 0.05 (0.69, 0.78)** | **0.73 ± 0.03 (0.70, 0.76)** | **0.73 ± 0.04 (0.70, 0.76)** |
| 2 | 0.68 ± 0.04 (0.65, 0.71) | 0.69 ± 0.05 (0.65, 0.73) | 0.65 ± 0.03 (0.62, 0.68) | 0.68 ± 0.04 (0.65, 0.71) | 0.68 ± 0.04 (0.65, 0.71) |
| 3 | 0.67 ± 0.04 (0.64, 0.71) | 0.66 ± 0.04 (0.63, 0.69) | 0.72 ± 0.07 (0.66, 0.78) | 0.68 ± 0.04 (0.64, 0.71) | 0.67 ± 0.04 (0.64, 0.71) |
| 4 | 0.69 ± 0.03 (0.67, 0.72) | 0.70 ± 0.03 (0.67, 0.73) | 0.67 ± 0.06 (0.63, 0.72) | 0.69 ± 0.03 (0.67, 0.72) | 0.69 ± 0.03 (0.67, 0.71) |
| 5 | 0.67 ± 0.02 (0.65, 0.69) | 0.66 ± 0.03 (0.64, 0.69) | 0.71 ± 0.03 (0.69, 0.74) | 0.67 ± 0.02 (0.65, 0.69) | 0.67 ± 0.02 (0.65, 0.69) |

Table 6: Inferred ChromAgeNet scores for aged HSC treated with compound.

| Condition | Mean | Stand. Dev. | Median | Count | 95% CI |
|---|---|---|---|---|---|
| Aged | 0.33 | 0.15 | 0.32 | 260 | (0.31, 0.35) |
| Aged + CASIN | 0.40 | 0.18 | 0.37 | 119 | (0.36, 0.43) |
| Aged + IOX | 0.56 | 0.14 | 0.55 | 31 | (0.51, 0.6) |
| Aged + RhoAi | 0.48 | 0.19 | 0.47 | 388 | (0.46, 0.5) |
| Aged + UNC | 0.55 | 0.16 | 0.52 | 46 | (0.5, 0.59) |
| Young | 0.56 | 0.19 | 0.54 | 109 | (0.52, 0.59) |

Table 7: Model hyperparameters for Grid Search

| Hyperparameter | LR | RF | XGBoost |
|---|---|---|---|
| C | [0.01, 0.1, 1, 10] | - | - |
| Penalty | ['l2', 'l1'] | - | - |
| N. Estimators | - | [100, 300, 500, 700, 900] | [100, 300, 500, 700, 900] |
| Max Tree Depth | - | [10, 20, 30, 40, 50] | [5, 10, 15, 20, 25] |

Table 8: Mu Fidelity metrics computed across many attribution methods over 1,000 confidently and correctly predicted images for each of the training classes.

| Attribution | Fidelity Young | Fidelity Aged |
|---|---|---|
| Saliency | 0.29 | 0.06 |
| DeconvNet | 0.01 | -0.04 |
| GradientInput | 0.03 | 0.01 |
| GuidedBackprop | -0.05 | 0.09 |
| IntegratedGradients | -0.17 | 0.00 |
| SmoothGrad | 0.03 | 0.00 |
| SquareGrad | 0.28 | 0.03 |
| VarGrad | 0.26 | 0.01 |
| GradCAM | 0.05 | 0.01 |
| GradCAMPP | 0.00 | 0.00 |
| Occlusion | -0.17 | 0.21 |
| Rise | 0.13 | 0.04 |
| SobolAttributionMethod | 0.27 | 0.01 |
| KernelShap | -0.21 | 0.06 |
| HsicAttributionMethod | 0.24 | -0.06 |
